# Human Genome-Scale Models of Metabolism and Gene Expression Reveal Resource Constraints of Cancer Cell Lines

**DOI:** 10.64898/2026.05.30.728988

**Authors:** Hratch Baghdassarian, Pablo Di Giusto, Juan Tibocha-Bonilla, Erick Armingol, Saratram Gopalakrishnan, Leo Dworkin, Laurence Yang, Nathan E. Lewis

## Abstract

Genome-scale metabolic models (M-models) provide mechanistic insight into intracellular metabolism by simulating fluxes subject to nutrient and energy resource constraints. However, they cannot account for a major component of resource allocation, since they do not explicitly account for the cost of producing and maintaining enzymes. Genome-scale models of metabolism and gene expression (ME-Models) address this by including gene expression reactions, but these have only been developed for prokaryotes due to the additional complexity and challenges of modeling eukaryotes. Here, we present the human ME-Model, which encodes transcription, translation, complex formation, and turnover reactions for all enzymes catalyzing metabolic reactions, and couples these processes to constrain metabolic fluxes. We introduce humanME, a Python package to build and analyze human ME-Models. With this, we constructed 16 cancer cell line ME-Models. We found that resource constraints improve growth-rate predictions, and that ME-Model flux predictions are more biologically plausible and efficient. Moreover, transcriptional fluxes recapitulate RNA-Seq expression levels, with discrepancies revealing potential trade-offs involving multiple cellular objectives. Finally, the ME-Model recapitulates the Warburg effect, with increasing growth rate inducing glycolytic shifts, in part due to machinery costs of the electron transport chain. Altogether, we show ME-modeling can mechanistically link gene expression, resource allocation, and metabolism in human cells, substantially expanding the predictive scope of constraint-based models.

## Introduction

A fundamental goal of systems biology is to quantitatively and accurately characterize how interactions between molecular components give rise to organism phenotypes and physiological functions^1^. Resource allocation provides a unique lens through which to view such genotype-phenotype relationships across multiple scales^2^. Under this theoretical framework, cell phenotype is subject to resource limitations, but also programmed to achieve a specific set of tasks (i.e., objectives) within said constraints^2,3^. Cells integrate environmental cues to decide and act upon biological objectives (e.g., growth^4^, secretion^5^, or cell migration^6^). Resource availability (e.g., nutrients, bioenergy, and macromolecular enzyme machinery) inform pathway activity such that cells can efficiently complete these objectives. When a cell encounters multiple objectives, the shared pool of limiting resources^7^ leads to trade-offs.

Genome-scale models (GEMs) of metabolism (M-Models) implement such an analysis framework by consolidating much biological knowledge and providing a context to integrate omics data and predict phenotypes^8^. Furthermore, since they maintain high-resolution molecular and mechanistic details, specific reaction fluxes can be studied. GEMs have provided key insights into wide-ranging biological systems, such as metabolic phenotypes underlying Alzheimer’s Disease^9^, polyamine metabolism in T-helper 17 cell pathogenicity^10^, cancer^11^, and countless other biomedical questions^12^. However, while M-Models explicitly account for nutrient and bioenergetic resources, they can only indirectly account for macromolecular machinery (e.g., via context extraction^13^ or flux minimization^14,15^).

A comprehensive accounting of macromolecular machinery costs is crucial not only for improving model accuracy, but also to extend the mechanistic details and potential applications of GEMs. This is because macromolecular synthesis costs represent a substantial portion of the overall cell resource budget^16^ and are demonstrated to affect cell activity^2^. For example, cells have evolved ribosomal features that maximize auto-catalytic activity^17^, demonstrate reduced expression of energetically costly proteins^5^, and alter their metabolic activity to minimize machinery costs^18^.

Recently, a variety of approaches have been developed to account for cell machinery costs, collectively called resource allocation models^19–21^. A prominent approach, genome-scale models of metabolism and expression (ME-Models)^22^, explicitly accounts for machinery costs by incorporating transcription and translation of reaction-catalyzing enzymes^23^ and coupling^24^ these enzymes to their respective metabolic reactions (Fig. 1). ME-Models encompass the entirety of the M-Model (the “metabolic module”). Thus, they are analyzed like an M-Model, but the additional machinery constraints resolve limitations such as constant biomass composition and large flux variability due to an underdetermined solution space^18^. Because ME-Models further encode transcription and translation reactions (the “expression module”), they uniquely enable variable RNA and protein biomass components and mechanistic insights to gene expression^25,26^. Furthermore, coupling of enzyme gene products to metabolic reactions explicitly accounts for machinery synthesis and maintenance costs, thus reducing the feasible solution space.

**Fig. 1:**
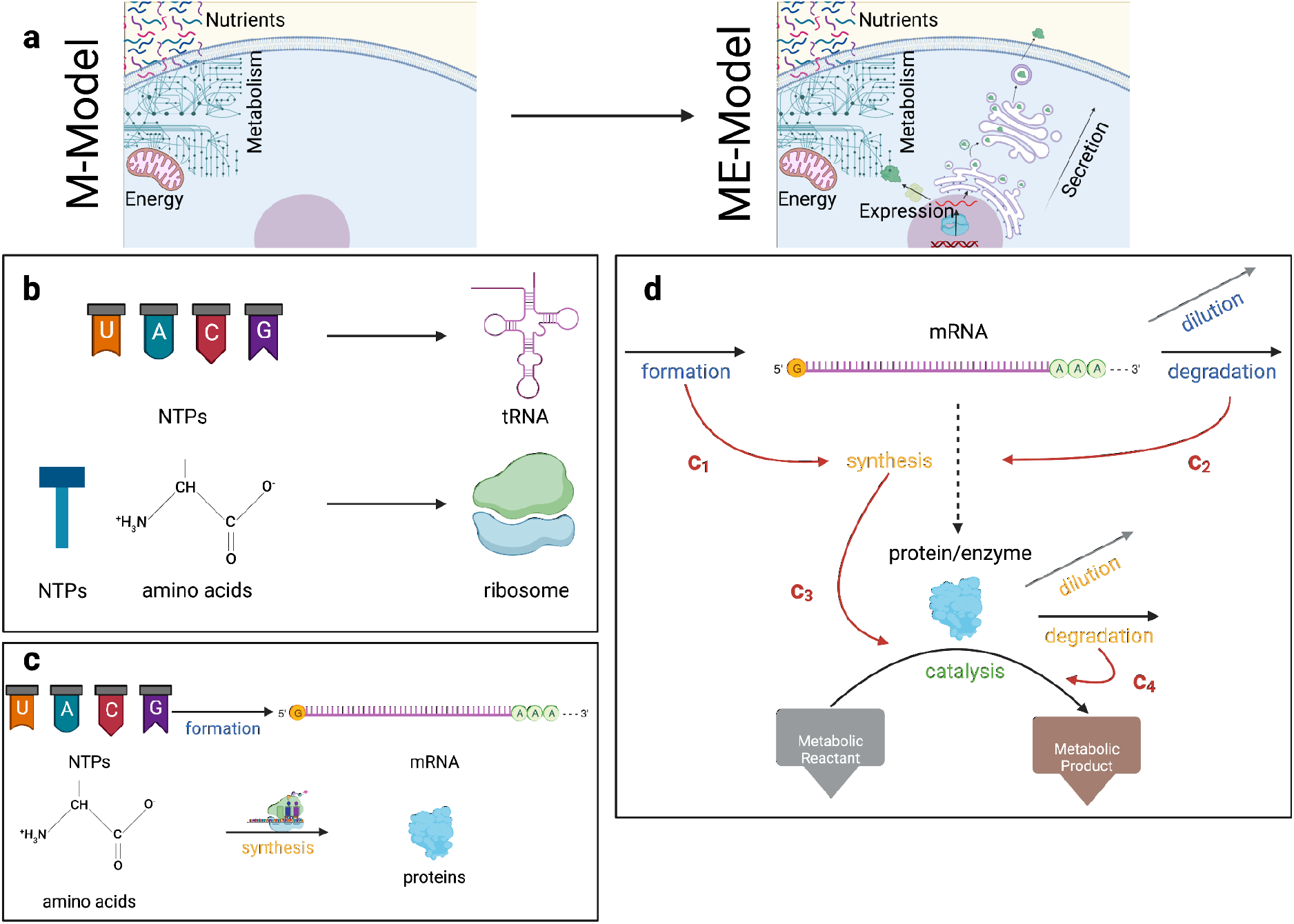
Steps for building a ME-Model. **(a)** An input M-model accounting for the metabolic module, as well as nutrient and energy resources, is used to construct a ME-Model, which can additionally account for machinery costs and optionally simulate the synthesis of non-metabolic proteins through the expression module. **(b)** tRNA and ribosomes are synthesized from metabolic precursors. **(c)** Gene-specific mRNAs and proteins are synthesized from metabolic precursors for each enzyme in the model. **(d)** A simplified reaction network of gene expression used to derive the coupling coefficients, i.e., parameters that link the different layers. Reactions for mRNA (blue text) are coupled to reactions for protein (orange text). Reactions for protein enzymes are coupled to the respective metabolic (or expression) reactions that they catalyze (green text). Black and gray arrows represent the gene expression reactions. The dashed arrow represents the fact that mRNA is not explicitly depleted by translation, but translation is dependent on mRNA concentration. The gray lines for dilution represent the fact that these reactions are not explicitly modeled for each macromolecule; rather, they are accounted for in the biomass dilution reaction. The red lines indicate which reactions are coupled to each other and are labeled by their respective coupling coefficient.

To date, ME-Models have not been built for eukaryotes, largely due to their additional complexities^27^, such as multiple subcompartments, relatively slower growth rates^28,29^, and non-growth objectives^18^. Here, we drew inspiration from a number of systems modeling approaches to create human ME Models: (1) ME-Models^22^ and similar resource balance analysis (RBA) approaches^30^ implemented in prokaryotes^31–34^, (2) GEMs that account for machinery costs in less explicit or more granular ways (e.g., parsimonious FBA^14,15^, thermodynamics-based constraints^35^, enzyme cost minimization^36^, GECKO^37^, constrained-allocation FBA^38^), (3) models that integrate multiple subsystems^39^, such as prokaryotic whole-cell models^1,40,41^ and mammalian GEMs of metabolism and the secretory pathway^5,18^, and (4) a formulation for unicellular eukaryotes by the yeast ETFL model^42^, which represents expression costs as primarily length-parameterized capacity constraints rather than explicit stoichiometric reactions, giving only indirect coupling to metabolic fluxes.

We present humanME, a Python tool built upon the COBRApy^43^ framework that constructs and analyzes cell-type or tissue-specific ME-models from a user-provided human metabolic model input. Like their prokaryotic counterparts, human ME-Models enable an exhaustive and direct accounting of machinery costs of metabolic activity. Beyond metabolism, they also simulate genome-wide transcriptional, translational, and enzyme transport fluxes. We first deployed humanME to build a ME-Model of the K-562 human immortalized myelogenous leukemia cell line. This ME-Model has improved growth rate predictions compared to a standard M-model. The additional constraints of the ME-Model also leads to more efficient and unique solutions for growth that eliminate thermodynamically infeasible loops but do not unnecessarily minimize flux through spontaneous reactions. Solutions also reduced flux variability analysis (FVA) flux ranges across key metabolic pathways with differing network topologies. Next, under a growth objective, we show agreement between FBA simulated transcriptional fluxes and RNA-sequencing measurements. A subset of unexpressed genes are associated with respiratory bioenergetic objectives that do not perfectly align with growth, implying multiple simultaneous cellular objectives. We also observe a strong Warburg effect-like preference for aerobic glycolysis over oxidative phosphorylation (OxPhos), with a monotonic glycolytic shift with growth rate. We attribute this Warburg effect in part to the OxPhos machinery costs, supporting the hypothesis that this phenomena occurs due to the relative proteome efficiency of generating flux through glycolysis compared to respiration^44^. Throughout these analyses, we observed limited tricarboxylic acid (TCA) flux, transcriptome distributions indicating competing growth and bioenergetic objectives, and the Warburg effect–which can be tied together via reductive glutamine metabolism to support lipid biosynthetic requirements of proliferation^45–47^. Finally, we construct an additional 16 ME-Models of various cancer cell lines, and demonstrate that, when nutrients availability is less constrained, ME-Model predicted growth rates are more accurate than their corresponding metabolic model.

## Results

### Building Human Genome-Scale Models of Metabolism and Gene Expression

The humanME software accepts an input context-extracted M-Model from Recon2.2^48^, and subsequently returns a corresponding ME-Model with coupled reaction-catalyzing enzymes (Fig. 1a). It does so in four steps as outlined below (for details, see Methods - Building the ME-Model). The tool also conducts analyses with the metabolic and expression modules. Due to the additional machinery constraints and small molecule demands imposed by the ME-Model, it is crucial to use an accurately curated M-Model as input to avoid feasibility issues when solving; in fact, the ME-Model can reveal key missing reactions that M-Model simulations overlook, resulting in more biologically plausible metabolic networks (see Supplementary Information and Results). The steps for generating the ME-Model are as follow:

#### Step 1

Generate expression reactions for ribosome biogenesis and tRNAs (Fig. 1b). While these are independent of the inputs, they are necessary for protein synthesis in the subsequent step (Fig. 1c).

#### Step 2

Generate gene-specific expression reactions (transcription, translation, and macromolecular degradation)^23^ for each catalytic protein, using the model’s gene-protein-reaction (GPR) rules (Fig. 1c). These proteins are also transported to the appropriate subcellular compartment where the reaction occurs, and form complexes according to the GPR. Since expression reactions themselves are catalyzed by enzymes (auto-catalytic), we recursively generate corresponding expression reactions until no new expression module enzymes are introduced. Bacterial ME-models assume protein degradation is negligible relative to growth rate, but this assumption does not hold in human cells.^28,29,49^ Thus, we include protein degradation.

#### Step 3

Couple each mRNA to its respective enzyme product^24^, and each enzyme to the metabolic reaction they catalyze, to generate flux demand throughout the expression module (Fig. 1d, Methods - Reaction Coupling).

#### Step 4

Re-formulate the M-Model biomass reaction to allow for variable RNA and protein biomass components (see Methods - Formatting the Biomass Reaction).

The final ME-models can then be subjected to various analyses, such as maximizing biomass, solving for non-growth objectives, and FVA.

While humanME provides consensus gene features necessary for Steps 1-3 (termed the protein-specific information matrix (PSIM)), users may optionally provide PSIMs to customize these features. Additionally, humanME can generate expression reactions for proteins that are not catalyzing any reactions in the M-Model, which enables exploration of such processes such as protein secretion, relevant to intercellular communication. Overall, our tool allows users to generate a context-specific human ME-Model for their cell type or tissue of interest.

### The ME-Model Metabolic Module Solution is More Efficient, Constrained, and Accurate

To assess the ME-Model, we used a M-Model of the K-562 erythroleukemic cell line^50^ as input (see Methods - Refining NCI-60 Cell Line M-Model Inputs). From the 2,554 reactions and 1,918 metabolites in the M-model, we obtained a ME-Model with 15,948 metabolites (1,911 metabolites and 1,778 expressed genes in various compartments and forms) and 60,022 reactions (6,052 metabolic reactions when separating isozymes and reversible reactions, and 53,952 gene expression reactions). While the M-Model predicted a growth rate of 0.056 hr^-1^, the additional machinery constraints implemented in the ME-Model reduced the predicted growth rate 0.025 hr^-1^. As compared to the experimental growth rate of 0.035 hr^-1^, the ME-Model prediction is more accurate than the M-Model prediction as quantified by the log_2_-fold-change (-0.47 and 0.66, respectively).

Next, we compared flux balance analysis (FBA) solutions between the ME-Model’s metabolic module and the M-Model to see if the additional machinery constraints yielded a more efficient solution, as quantified by the total absolute flux through all metabolic reactions (Fig. 2a). We expected that the additional biosynthetic costs of gene expression would enforce more efficient use of the ME-Model to achieve its objective. This is analogous to parsimonious flux balance analysis (pFBA)^14,15^ and the Max-Min Driving Force (MDF)^35^. To make this comparison fair, we assessed the efficiency of both models at the same growth rate (the maximum of the ME-Model, 0.025 hr^-1^). The ME-Model predicts a total flux approximately four orders of magnitude lower than the M-Model (25.3 vs 37.2e4 mmol/gDWcell/hr), significantly smaller than the sampled flux distribution (p << 1e-3). We reasoned that this difference is due, in part, to the elimination of thermodynamically infeasible cycles (type III pathways or “loops”^51^). Thus, we ran the M-Model FBA solution through the CycleFreeFlux algorithm^52^ to eliminate thermodynamically infeasible loops from this solution. This resulted in a much more comparable efficiency. Interestingly, the efficiency of the ME-Model lies between that of pFBA (15.6 mmol/gDW_cell_/hr) and CycleFreeFlux (36.1 mmol/gDW_cell_/hr). This indicates that beyond eliminating type III loops, the ME-Model also minimizes machinery costs (e.g., penalizing type II futile cycles).

**Fig. 2:**
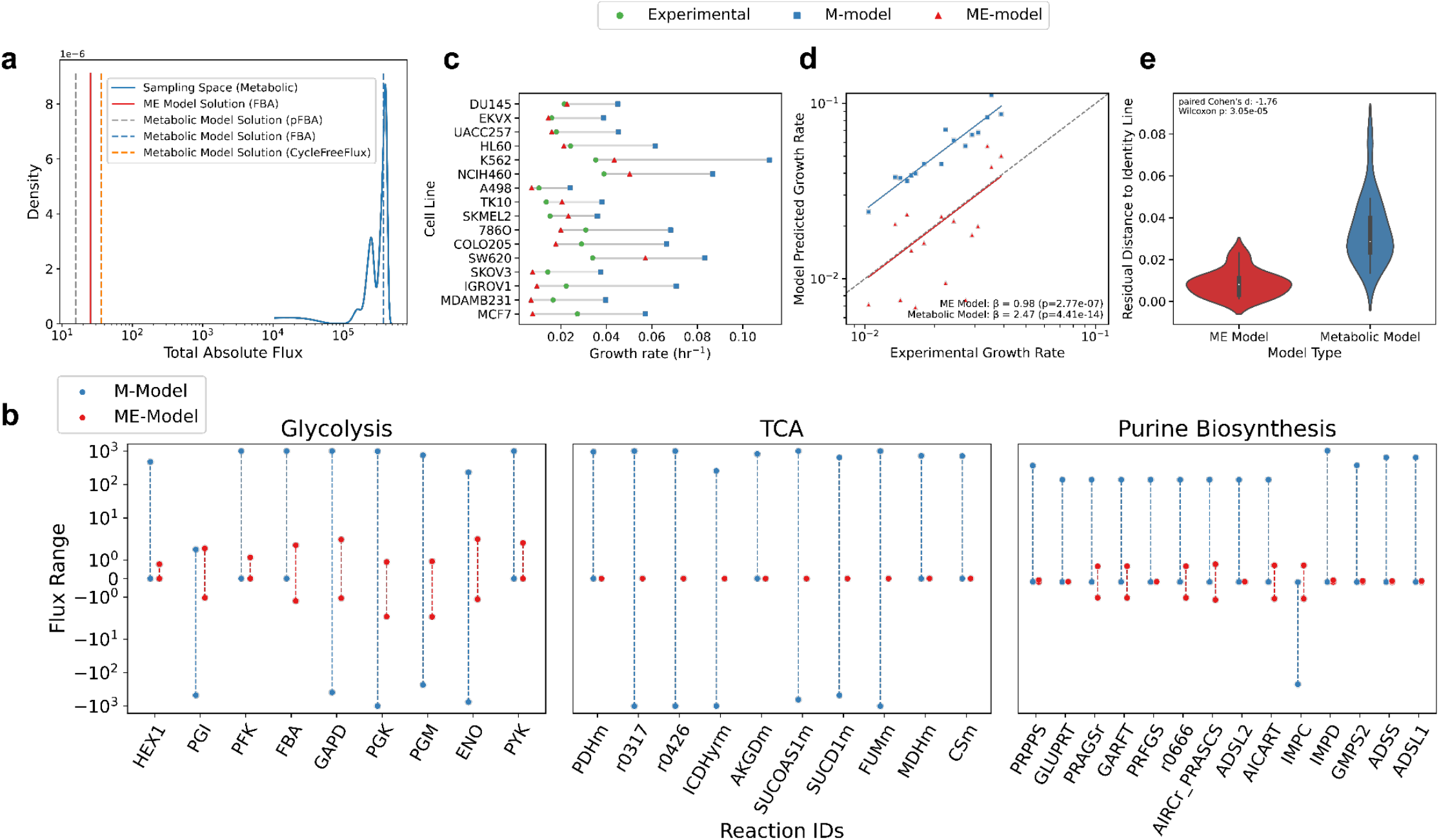
Comparison of metabolic flux from ME-Model’s metabolic module and the M-Model. **(a)** FBA solution efficiency quantified by the total absolute flux through all metabolic reactions (x-axis). All FBA was run at the maximum ME-model growth rate. The kernel density estimate (y-axis) of the M-model solution space obtained from 1000 samples is shown in blue. Vertical lines indicate the M-model FBA solution (blue dashed), CycleFreeFlux50-corrected M-model FBA solution (orange dashed), ME-model FBA solution (red), and parsimonious FBA (pFBA) solution of the M-model (gray dashed). **(b)** FVA of the ME-Model (red) and the M-Model (blue) for reactions in glycolysis (left panel), the TCA cycle (middle panel), and purine biosynthesis (right panel). Individual reactions (x-axis) are plotted against minimum and maximum fluxes (y-axis, symlog scale) that maintain 95% of the maximal growth rate identified by each model. **(c)** Experimental growth rates alongside M- and ME-Model maximal growth rates across the 16 cell lines for which feasible matched ME-Model/M-model solutions were obtained. FBA simulations are run with exchange reaction bounds relaxed by 2-fold. Cell lines are ordered by the most accurate ME-Model predictions. The ME-Model systematically predicts lower maximal growth rates than the paired M-model, consistent with the additional machinery constraints imposed by the expression module. **(d)** Scatter plots and OLS regression lines compare model predicted growth rates to experimental growth rates, with axes subsequently plotted on a log-log scale. OLS coefficients are annotated alongside their Wald test p-values. Coefficients were estimated using models without an intercept to enable direct comparison to the identity line y=x (gray), with coefficients closer to 1 indicating more accurate predictions across cell lines. **(e)** Violin plots summarize the absolute distance of each scatter point in panel (d) to the identity line, representing deviation from perfect prediction.

Given that the underdetermined nature of M-Models^18^ results in a large solution space^53^, we tested if the explicit accounting of machinery resource costs would identify a more constrained solution. Thus, we conducted FVA on the M- and ME-Models at 95% of their respective maximum growth rates. Due to the computational demand of solving the ME-Model, we ran FVA for reactions from three selected pathways representing different network topologies: a linear pathway (glycolysis), a cyclic pathway (the TCA cycle), and a branching pathway (purine biosynthesis) (Fig. S1a). Comparing the flux ranges between the two models, the ME-Model is substantially more constrained than the M-Model across all reactions (Fig. 2b). Notably, ME-Model flux ranges in glycolysis were nearly three orders of magnitude larger than those in the TCA cycle (Fig. S1b), based on the 90th-percentile flux range across reactions (log_10_ ratio = 2.94). This result is directionally consistent, though larger in magnitude, with the well-established dominance of glycolytic throughput over oxidative metabolism in proliferative cells, where glycolytic fluxes can exceed TCA cycle fluxes by up to two orders of magnitude during the Warburg effect^54^.

To examine whether accounting for proteome constraints generalized beyond K-562, we next screened a subset of sixteen additional NCI-60 cancer cell lines for which context-specific M-models were available(Fig. 2c). While only 6 of 16 ME-Models had more accurate predictions quantified by log-fold-change (LFC) relative to experimental growth rates, we found no significant differences in global predicted performance between the two model types (Fig. S4c; RMSE of 0.01 for both model types). Metabolic models tended to over-predict growth whereas ME-Models tended to under-predict growth at similar levels (OLS coefficients relative to β = 1; Fig. S4a-b), consistent with the additional machinery constraints imposed by the expression module.

Growth rate under-prediction could reflect genuine intracellular resource-allocation constraints or overly restrictive extracellular metabolite exchange bounds inherited from the M-Models. To distinguish between these possibilities, we relaxed exchange bounds 2-fold across cell lines. This can reflect contexts in which extracellular nutrient availability is not a limiting factor on cell growth. Consequently, under conditions in which extracellular nutrient limitation was minimized, ME-model growth predictions were globally more accurate than those of the corresponding metabolic models (RMSE of 0.01 vs 0.04; 14 of 16 ME-Models have smaller relative deviation in prediction from experimental growth by LFC; Fig. 2c–e). This indicates that explicitly accounting for proteome constraints improves predictive fidelity when growth is not dominated by nutrient availability. Overall, the growth-limiting effect of explicit machinery accounting generalizes across multiple cellular contexts, while also highlighting that model feasibility and quantitative calibration remain important bottlenecks for broader deployment of human ME-Models.

Together, these results demonstrate that the ME-Model does not merely constrain flux magnitude, but reshapes the feasible solution space in a mechanistically relevant manner by coupling metabolism to the costs of gene expression.

### The K-562 ME-Model Exhibits Multiple Competing Objectives and the Warburg Effect

Unlike standard M-Models, ME-Models explicitly represent gene expression flux. We asked whether first-principle–derived coupling constraints and flux-balance–based resource allocation alone are sufficient to recover accurate relative transcriptional fluxes at the genome scale. Under a growth objective constrained to 95% of maximal growth, K-562 ME-Model transcriptional fluxes showed substantial rank agreement with measured transcript abundances across 1,765 genes (Spearman ρ = 0.36), despite transcript levels not being directly constrained by or calibrated to expression data (Fig. 3a). This agreement was not driven by zero-valued transcriptional fluxes (non-zero flux Spearman ρ = 0.33) and substantially exceeded correlations expected by chance (one-sided permutation p = 9.99e-5).

**Fig. 3:**
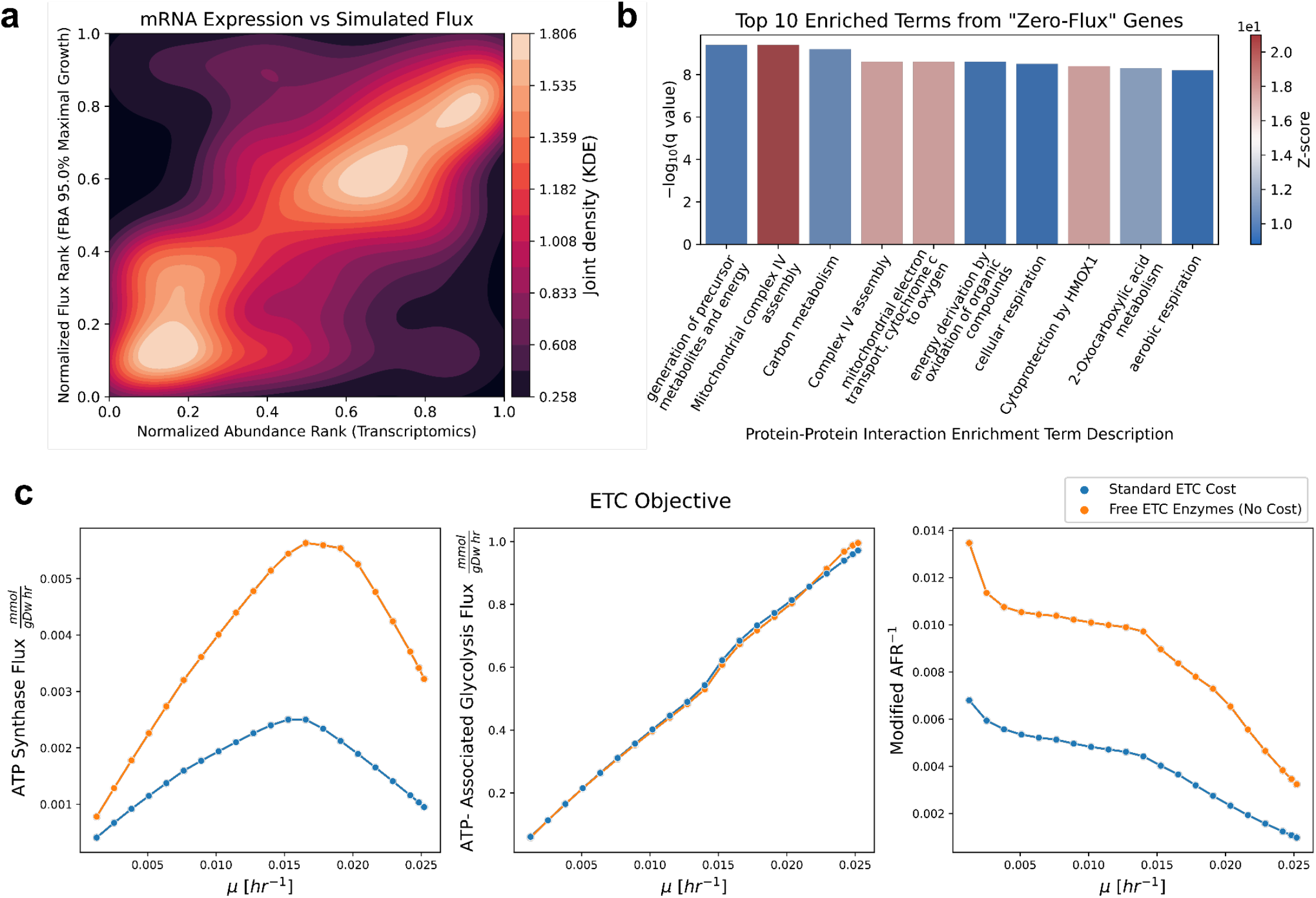
ME-Model Proteome Constraints Reveal Coordinated Expression and Metabolic Changes Associated with the Warburg Effect. **(a)** Two-dimensional kernel density estimate (KDE) plot illustrating the joint distribution of normalized transcriptomic abundance ranks (x-axis) and normalized transcriptional flux ranks (y-axis) obtained from flux balance analysis (FBA) at 95% of maximal growth, with growth as the objective. Each axis represents per-feature ranks scaled between 0 and 1. The KDE contours depict smoothed probability density estimated over the bivariate rank space. The color bar indicates the estimated joint probability density of observing particular combinations of normalized transcript abundance and flux ranks. Brighter colors correspond to higher densities, highlighting regions where transcript and flux ranks co-occur more frequently. **(b)** Bar plots of the top 10 enriched terms for growth objective “zero-flux” genes from Metascape’s protein–protein interaction enrichment analysis, ordered by Benjamini–Hochberg FDR-corrected q-values and colored by z-score. **(c)** Total pathway fluxes as a function of growth rate with the electron transport chain (ETC) set as the objective, shown for ATP-associated reactions in oxidative phosphorylation (OxPhos) and glycolysis (left and middle panels). Pathway fluxes were aggregated by subtracting the summed ATP-consuming reactions from summed ATP-producing reactions, yielding the inverse modified AFR (i.e., inverse of the glycolysis to OxPhos ratio) shown in the right panel. Scatter points represent individual FBA solutions, while lines connect solutions across growth rates. Blue lines represent the solutions with standard ME-Model formulation, and orange lines represent those with removed proteome costs for ETC enzymes.

Next, we sought to identify the sources of discrepancy between ME-Model transcriptional fluxes and measured mRNA abundances. We first asked whether ‘OR’ GPR isozymes, which generate functionally redundant ME-Model reactions, lead the model to preferentially select the less resource-costly alternatives. Thus, we repeated the correlation analysis after restricting isozymes to those with the maximal transcriptional flux. This reduced the correlation (Spearman ρ = 0.21), suggesting that isozyme redundancy is unlikely to be the primary source of the discrepancy. Because growth is not always the most accurate objective in mammalian cells^2,18^, we tested whether alternative objectives or metabolic hedging could better explain the discrepancy^55^. Specifically, we observed 210 “zero-flux” genes that had substantial expression (log_2_(TPM + 1) ≥ 2) but no transcriptional flux (Fig. S2a-b), potentially reflecting non-growth-associated objectives. Enrichment analysis of these genes revealed strong overrepresentation of electron transport chain (ETC)–related processes among the top ten protein–protein interaction–based terms (Fig. 3b). Consistent with this signal, FBA simulations revealed incomplete transcriptional synthesis of key ETC components. For Complex II, only a subset of required monomeric subunits exhibited nonzero transcriptional flux (⅓ and ½ of subunits for each isozyme), resulting in no corresponding metabolic flux, despite the reaction being feasible and unblocked. More broadly, across Complexes I–IV, monomeric subunits exhibited progressively reduced transcriptional synthesis flux relative to other genes in the model as growth rate increased (Fig. S2c). While energy generation supports biomass production, these processes are not directly coupled^56^, and bioenergetic pathways can trade-off with growth, drawing carbon flux away from biomass synthesis. This effect is particularly pronounced for respiratory pathways under high proliferation, where aerobic glycolysis is often favored over respiration (the Warburg effect). Furthermore, enrichment of “Cytoprotection by HMOX1” suggests hedging against future stress conditions, including hypoxia. An extended discussion of the interplay between growth, energy production, and hedging is provided in Appendix A and Appendix D.2 of Baghdassarian *et al*.^*2*^

To evaluate ETC as an alternate objective to growth, we analyzed flux changes when the ME-Model objective is set to maximize ETC activity. The maximal ETC flux was 0.0025 mmol/gDw/hr and occurred at a growth rate of 0.015 hr^-1^ (Fig. S3b, left panel). While the correlation between transcriptional flux and mRNA abundance remained unchanged (Spearman ρ = 0.36 at 99% of maximal ETC, Fig. S2d), transcriptional fluxes under ETC and growth objectives were not perfectly correlated (Spearman ρ = 0.80), indicating partial reallocation of expressed genes between objectives. Specifically, 470 (26.6%) of the 1,765 genes exhibited substantial rank shifts (Fig. S2e). The sets of “zero-flux” genes exhibited moderate overlap between growth and ETC objectives (Jaccard index = 0.31). When enrichment analysis of “zero-flux” genes was repeated for the ETC objective, concordance at the pathway level decreased further, with a Jaccard index of 0.27 across all significant pathways (FDR ≤ 0.1; 592 and 904 significant pathways for growth and ETC objectives, respectively). Notably, the top ten enriched pathways remain associated with ETC-related terms (Fig. S2f), indicating that despite global proteome reallocation, the dominant enrichment signal reflects persistent absence of transcriptional flux in ETC-associated genes rather than objective-specific upregulation—consistent with a high proteome cost of respiratory metabolism.

Consistent with this interpretation, maximizing the ETC objective revealed a concave-down parabolic relationship between ETC flux and growth rate (Fig. S3c, left panel). This suggests that cells shift away from respiratory ATP production as growth increases, motivating further examination of the Warburg effect in our simulations. To quantify this, we used the ATP flux ratio (AFR), defined as the ratio of mean glycolytic to oxidative phosphorylation (OxPhos) ATP-producing reaction fluxes. This metric has been previously applied to M-Models of NCI-60 cancer cell lines^57^. We modified the AFR to account for ATP-consuming reactions (Fig. 3c blue line, S3a; see Methods); results were qualitatively unchanged when using the original AFR^57^ (Fig. S3b) or total ETC and glycolysis fluxes (Fig. S3c). The AFR reveals several consistent trends: (1) OxPhos is a concave-down parabolic function of growth, indicating that while respiratory energy generation is initially coupled to biomass production, the two objectives diverge beyond the inflection point of 0.015 hr^-1^. This behavior is consistent with Warburg-like trade-offs observed at high proliferation rates. (2) Flux through glycolysis increases more rapidly than that of OxPhos even below this inflection point, resulting in a monotonic shift toward glycolytic flux across all growth rates. This is consistent with evidence that the Warburg effect occurs even at low growth rates^56^, and consistent with the notion of hedging for future hypoxic conditions. (3) Glycolytic flux exceeded OxPhos across all growth rates by more than two orders of magnitude (median log_10_(modified AFR) = 2.35; IQR 2.29–2.63), consistent with observations of glycolysis contributing a substantial fraction of ATP production during the Warburg effect (up to 64%^58^). Order-of-magnitude differences are consistent with prior descriptions of Warburg phenotypes, in which glycolytic flux exceeded TCA cycle flux by up to ∼100-fold^54^.

### ETC Proteome Costs Contribute to Warburg-Like Metabolism in the K-562 ME-Model

Proteome efficiency has been proposed to drive aerobic glycolysis at high proliferation rates^44^: when normalized by enzyme molecular weight, glycolysis can yield more ATP per unit proteome investment than respiration. The coupling constraints implemented in the ME Model explicitly impose proteome costs. Given our observations of preferential glycolytic flux (Fig. 3c, right panel) and the reduced investment in ETC enzyme production with increasing growth rate (Fig. S2c), we tested the proteome-efficiency hypothesis by removing the proteome cost of ETC proteins. Upon confirming that this trend is not from nutrient limitation (Fig. S3d), we re-evaluated the pathway fluxes after allowing the ME-Model to freely synthesize Complexes I-IV (by adding sink reactions; Fig. 3c, green line). Without respiratory complex enzyme costs, the ME-Model achieved a maximal ETC flux of 0.0056 mmol/gDw/hr, more than doubling the value obtained under the standard formulation (Fig. 3c, blue line). Notably, the concave-down parabolic relationship between ETC flux and growth rate remained qualitatively similar, with ETC flux maximized at a growth rate of 0.016 hr^-1^. However, total ETC fluxes shifted to higher values across all growth rates following removal of ETC proteome costs. In contrast, glycolytic flux was unchanged, resulting in a significant shift toward respiration across growth rates (one-sided paired Wilcoxon signed-rank test, *p* = 4.8e-7, paired Cohen’s *d* = 3.97, median free-to-costly log_2_ modified AFR^-1^ ratio of 1.135 (IQR [1.017, 1.491]) across growth rates). Altogether, these results indicate that ETC protein costs directly contribute to the Warburg effect; however, given that the parabolic relationship between respiratory flux and growth rate is preserved even when ETC enzymes are free, additional reactions and enzymes in the network likely constrain respiration, preserving the overall glycolytic preference of the ME-Model.

### K-562 ME-Model Simulations Exhibit Lipogenic Flux

When conducting FVA to characterize the ME-Model solution space (Fig. 2b; Fig. S1b), we observed that reactions supporting citrate production—specifically citrate synthase (“CSm”) and mitochondrial isocitrate dehydrogenase (“ICDHyrm”)—exhibited larger maximum values and stronger directional preferences than the rest of TCA (Fig. S1b). Citrate and acetyl-CoA are key metabolic precursors for lipogenesis, which supports biomass generation for growth (Fig. 3a). CSm generates citrate from substrates oxaloacetate (OAA) and acetyl-CoA in the canonical TCA direction, whereas ICDHyrm exhibited a preference to proceed in the reverse direction, reductively carboxylating α-ketoglutarate (αKG) to produce isocitrate.

These patterns implicate both glucose and glutamine-derived carbon in support of lipogenesis: CSm substrates can be derived from either glucose or glutamine carbon, while reverse ICDHyrm flux specifically implicates glutamine. Glucose-derived pyruvate can contribute to OAA production via pyruvate carboxylase and acetyl-CoA production via pyruvate dehydrogenase. Glucose-derived carbon is demonstrated to direct flux toward citrate production via the TCA cycle during the Warburg effect^46,59^.

Additionally, a recurring metabolic attribute associated with the Warburg effect is the use of glutamine-derived carbon to support both TCA intermediate anaplerosis and lipogenesis for biomass generation^60^. Glutamine-derived αKG can support lipogenic citrate production via two routes: through reductive carboxylation as described above or oxidatively. In the oxidative direction^46^, αKG proceeds through the canonical TCA cycle to malate, which can either (i) be converted to OAA by malate dehydrogenase and condense with acetyl-CoA at citrate synthase, or (ii) be decarboxylated to pyruvate by malic enzyme, generating NADPH that supports the reductive steps of fatty acid synthesis. Glutamine-derived pyruvate is typically fermented to lactate by lactate dehydrogenase to maintain redox balance. Reductive carboxylation has been shown to be favored under hypoxic and normoxic conditions at high proliferation rates^45^, and in cells with suppressed ETC activity^47^. The pathway is even used in quiescent fibroblasts^61^. Together, these prior studies provide a framework for interpreting the elevated CSm and reverse-ICDHyrm flux observed in our ME-Model: the Warburg effect rewires not only glycolysis and bioenergetics, but also glutamine metabolism to support lipogenic citrate production.

To explore the extent to which glutamine- and glucose-derived carbon was diverted to support lipogenesis, we visualized the FBA and FVA results of the associated reactions across growth rates (Fig. 4b; Fig. S5). We found that glucose-derived pyruvate could undergo lactic acid fermentation or be converted to acetyl-CoA to fuel CSm, whereas pyruvate carboxylase was pruned during context extraction. Glutamine-derived αKG could proceed either through mitochondrial reductive carboxylation or through oxidative TCA reactions, with both malic enzyme and cytosolic reductive carboxylation absent from the context-extracted K-562 model. However, we observed a slight directional preference towards reductive carboxylation for isocitrate dehydrogenase in FVA. In contrast, the canonical-direction oxidative TCA reactions downstream of αKG showed comparatively weak directional preference; though, these reactions remained available and carried the forward flux needed to supply OAA for CSm, given that pyruvate carboxylase was pruned. Lipogenesis reactions downstream of citrate exhibited positive flux preferences, indicating production of lipids. Altogether, these results indicate that both glucose- and glutamine-derived carbon are preferentially converted to citrate for lipogenesis: glucose carbon enters as acetyl-CoA via PDH, while glutamine carbon contributes through both reductive and oxidative mitochondrial routes, rather than exclusively through reductive carboxylation.

**Fig. 4:**
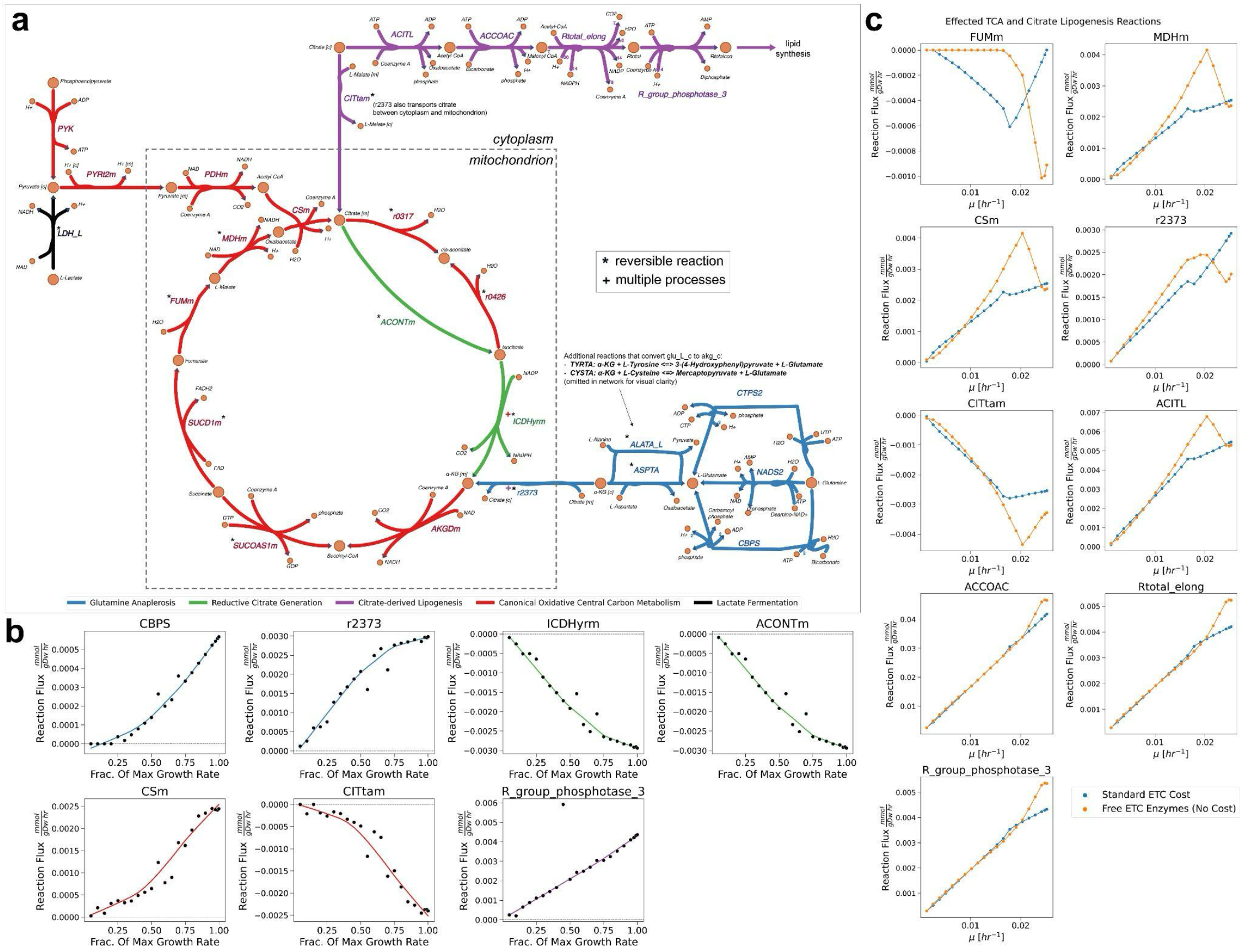
Metabolic flux rewiring across growth rates to support lipogenesis. **(a)** Reaction network of the TCA cycle, glutamate catabolism, and lipogenesis (colored by process, with a “+” if associated with a second process). Reaction arrows point in the forward direction as specified by Recon2.2, with reversible reactions denoted by an asterisk. ICDHyrm in the forward direction is participating in reductive citrate generation and canonical oxidative central carbon metabolism in the reverse and forward directions; r2373 mediated mitochondrial import of alpha-αKG and export of cytosolic citrate in the forward direction is associated with glutamine anaplerosis and citrate-derived lipogenesis, respectively. **(b)** Individual reaction fluxes as a function of growth rate identified by FBA under a growth objective. Key reactions from various metabolic processes highlight diversion of flux towards citrate lipogenesis, colored as in panel (a). The x-axis is normalized to the maximum ME-Model growth. Scatter points visualize individual FBA solutions, whereas lines visualize loess regression curves. **(c)** Individual reaction fluxes as a function of growth rate identified by FBA under an ETC objective. Scatter points represent individual FBA solutions, while lines connect solutions across growth rates. Blue lines represent the solutions with standard ME-Model formulation, and orange lines represent those with removed proteome costs for ETC enzymes. All fluxes associated with TCA and citrate lipogenesis were explored, but only those that showed differences between the two solutions are displayed.

### ETC Proteome Costs Rewire TCA Activity

Metabolic alterations to glutamine and glucose metabolism, as well as TCA cycle flux, are well-documented features of the Warburg effect. Yet, to our knowledge, no study has causally linked lipogenesis, TCA flux, and ETC activity. The connection has two possible directions: (1) diverting flux toward citrate production limits the oxidative TCA flux necessary to produce the reducing cofactors used by the ETC, as articulated by Yu *et al*.^*62*^ or (2) impaired or limited ETC activity promotes a rewiring of TCA metabolism, favoring non-oxidative TCA metabolism and reductive carboxylation, as articulated by Mullen *et al*.^47^ In this second context, the proteome costs imposed by the ME-Model can be viewed as encoding endogenous constraints that effectively limit ETC capacity; consequently, removing ETC proteome costs tests whether the observed citrate-associated rewiring is downstream of respiratory capacity rather than upstream cofactor supply. To test the first direction, we provided an unlimited supply of mitochondrial NADH and FADH_2_ (by adding sink reactions for them); we observed no change to total ETC flux or the modified AFR, indicating that this pool of reduced mitochondrial cofactors does is not the dominant limitation on ETC in the ME Model. For the second direction, we next examined the effect of removing the proteome cost of ETC proteins (as described previously; Fig. 3c)– effectively decreasing ETC limitation--on TCA and citrate lipogenesis. Visualizing the reactions that had a noticeable difference in FBA solution flux (Fig. 4c), we see that canonical oxidative TCA flux is not simply restored. Instead, metabolic rewiring intensifies the lipogenic citrate phenotype rather than reversing it, with changes localized to specific reactions at higher growth rates. Malate dehydrogenase (MDH) flux increases, supplying additional OAA for citrate synthase; this may be mechanistically enabled by the relief of ETC proteome cost, since MDH’s forward direction generates NADH that can be cleared by increased ETC activity. The remaining oxidative TCA reactions remain inactive, and the additional malate consumed by MDH is supplied via the citrate–malate antiporter (“CITtam”). Together, increased MDH activity (and downstream citrate synthase activity) and enhanced citrate–malate exchange elevate citrate production and downstream lipogenesis. Notably, many of these changes occur at the parabolic inflection point growth rate for OxPhos flux (Fig. 3c), consistent with ETC capacity acting as the binding constraint on this rewiring. These results suggest that ETC capacity gates lipogenic citrate output through redox-mediated control of MDH flux and OAA supply.

Altogether, these results indicate that proteome allocation to respiratory complexes is a major constraint on ETC flux, and that relieving this constraint triggers a coordinated reorganization of mitochondrial metabolism— amplifying both respiratory and citrate-associated lipogenic fluxes—without fully restoring canonical oxidative TCA flux under the remaining model constraints.

## Discussion

In this work, we present a human genome-scale ME-Model that explicitly links metabolism, gene expression, and proteome allocation, enabling systematic exploration of how cellular resource constraints shape feasible metabolic states. We provide a Python tool, humanME, that enables users to build ME-Models from input context-specific human metabolic models. We demonstrate that the additional machinery constraints imposed in the ME-Model capture a biologically grounded, efficient solution that eliminates thermodynamically infeasible loops and reduces the solution space. A unique feature of ME-Models relative to M-Models is that they explicitly simulate gene expression fluxes, enabling direct comparison to measured transcript abundances without calibrating the model to expression data. We found that ME-Model transcriptional fluxes recapitulated RNA-Seq relative abundance, supporting the validity of first-principles coupling constraints for predicting genome-wide expression patterns. Importantly, the discrepancies between simulated flux and measured abundance were themselves informative. A set of highly expressed genes that carried no simulated transcriptional flux under a growth objective were strongly enriched for ETC components. This pattern suggested that the cell expresses respiratory machinery for purposes not fully captured by a growth objective alone, motivating us to examine competing bioenergetic objectives and, in turn, the Warburg effect.

Motivated by this observation, we examined whether the model recapitulated the Warburg effect. We observed that a Warburg-like effect emerged without imposing explicit flux bound constraints on glycolytic or respiratory pathways, characterized by a monotonic shift toward glycolytic flux across growth rates and a concave-down relationship between respiratory flux and proliferation. These observations align with previous work in GEMs demonstrating that while stoichiometric constraints alone are not sufficient to explain the Warburg effect, accounting for enzyme constraints can recapitulate this metabolic phenotype^63^. Perturbation experiments further demonstrate that this phenotype is not driven by cofactor scarcity from TCA rewiring, but instead by allocation of proteomic resources to respiratory machinery. Removing ETC proteome costs substantially increases ETC flux and attenuates the Warburg effect. Importantly, relieving this constraint induces coordinated rewiring of mitochondrial metabolism, increasing flux through malate dehydrogenase, citrate synthase, and downstream lipogenic pathways without re-establishing a fully oxidative TCA cycle. Together, these results indicate that global proteome allocation sets respiratory capacity, and mitochondrial carbon routing adapts accordingly to balance bioenergetic and biosynthetic demands. More broadly, this work illustrates how ME-Models can predict fluxes and mechanistically interrogate how cellular objectives, resource limitations, and network topology give rise to emergent metabolic phenotypes.

The ME-Model has limitations to be addressed in future work. The stringent machinery constraints that enable mechanistic insight also limit model feasibility or ability to simulate physiologically relevant growth rates; while this is useful for identifying missing essential reactions that corresponding metabolic models may bypass, systematic reaction curation remains a bottleneck. In constructing multiple ME-Models, we addressed this by relaxing nutrient uptake constraints. Although this reflects a biologically relevant context in which nutrient limitation is not a strong constraint, it reduces consistency with experimentally-constrained conditions (e.g., exometabolomics) that metabolic models can more readily accommodate. Furthermore, limitations in a metabolic model will also influence performance of its corresponding ME-Model. For example, these models do not incorporate regulatory and signaling networks^39^ and, when using FBA, they do not account for dynamic contexts in which metabolite concentrations are not at steady-state. Furthermore, in developing the expression module, assumptions and simplifications were made. Examples include deterministic gene expression without a probabilistic component to account for stochasticity, 1:1 stoichiometric ratio of enzyme complex subunits, the use of a dummy protein for orphan reactions, and estimation of enzyme transport costs as a function of protein length. Such assumptions should be refined to improve simulation accuracy. Additionally, the coupling constraints that link the expression module to the metabolic module require extensive knowledge of kinetic parameters, which are often imputed when unknown. Finally, because the ME-Model adds many reactions, computational time for solving becomes a bottleneck, particularly when trying to run model-wide analyses such as FVA of all reactions.

The ME-model offers several capabilities and future applications that warrant deeper exploration. These include identifying and refining kinetic parameters from flux simulations^64^, extending the framework to account for non-steady-state conditions^65,66^, and improving poorly characterized regions of the network. Such network refinements are particularly relevant for components of the expression module, including proteostasis networks, as well as for under-resolved areas of the metabolic module, such as lipid metabolism. Furthermore, a fundamental yet unexplored capability of the tool is to produce user-specified non-machinery proteins in any cell compartment of interest, including secretion of proteins to the extracellular matrix. As such, the model can identify therapeutic production bottlenecks and account for non-metabolic objectives such as cell-cell communication and migration (e.g., via proxy ATP-consuming motility reactions catalyzed by relevant machinery).

Altogether, the ME-Model represents an advance towards whole-cell modeling in mammalian systems, enabling detailed mechanistic interrogation of fundamental biological relationships, such as growth and bioenergetics. Beyond metabolism, this framework provides a foundation for exploring how gene expression constraints give rise to diverse phenotypes—including, but not limited to, growth^4^—across cellular contexts. By explicitly encoding a broad range of cellular processes, the ME-Model may also serve as a building block for multicellular tissue-scale modeling. For example, we can adapt approaches developed for tissue-specific^67^ and multicellular M-Models^68,69^ and integrate with complementary modeling frameworks for cell-cell communication^70^ and signaling^39^ through hybrid modeling strategies^71^. Together, this work positions the ME-Model as a foundation for mechanistic, multi-scale studies of how gene expression and metabolic constraints jointly shape cellular and, ultimately, tissue-level phenotypes.

## Methods

### Building and Solving the ME-Model

We constructed genome-scale human metabolism and expression (ME) models by augmenting input context-specific Recon2.2 metabolic networks with explicit gene expression, macromolecular synthesis, transport, degradation, and coupling constraints. Central to model construction is a gold-standard Protein-Specific Information Matrix (PSIM), which encodes gene- and isoform-specific sequence features, degradation rates, protein–RNA ratios, and localization information required to parameterize transcription, translation, folding, transport, complex formation, and degradation reactions. Isoform sequences were obtained from MANE Select^72^, RefSeq Select^73^, or APPRIS^74^, and missing features were filled using curated defaults. Gene expression is modeled across the full central dogma, encompassing pre-mRNA transcription, processing (capping, polyadenylation, splicing, and export), translation, protein folding, and compartment-specific transport and degradation via pathways including the ubiquitin-proteasome system, lysosomal degradation, and organelle-specific proteases. Metabolic and expression layers are linked through theoretically derived coupling coefficients that relate mRNA and protein fluxes to catalytic reaction fluxes under steady-state assumptions, incorporating gene-specific parameters such as protein-to-RNA ratios, degradation rates, and estimated catalytic rates. The biomass reaction is reformulated to allow variable RNA and protein mass fractions, with expression reactions directly contributing to biomass accumulation. Growth rate maximization is achieved through a binary search algorithm using the high-precision qMINOS solver^75^, which accommodates the large range of stoichiometric and coupling coefficient magnitudes inherent to ME-Models. Objectives other than growth were optimized by scanning across feasible growth rates Full details of the building and solving procedure are provided in the Supplementary Methods.

### Refining NCI-60 Cell Line M-Model Inputs

Input M-models were adapted from those generated in Richelle et al.^50^. Specifically, we took the mCADRE^76^-extracted M-models (not “Protected”) and modified the biomass reaction and the exchange reaction bounds. We also added missing reactions preventing model feasibility. We briefly describe some of these modifications below. Additional details regarding the biomass objective and missing reactions can be found in the Supplementary Information and Results.

The biomass reaction was re-formatted from the net reaction to reactions for formation of each biomass component separately, matching the format of Recon 2.2. Richelle et al.^50^ calculated the biomass reaction stoichiometric coefficients using “Table S1” from the “Supplementary Information” of ref^13^. Here, we combined the “resolved” and “unresolved” lipid mass fractions, representing mass fractions of 7.98e-2 and 2.27e-2, respectively, into a single lipid component by taking the sum (lipid mass fraction = 1.025e-1). Additionally, ATP hydrolysis for growth-associated maintenance (GAM) (“Growth-associated” and “Unresolved other components” from “Table S1”^13^.) have a mass fraction of 4.37e-2. However, GAM ATP hydrolysis should not have a direct mass fraction. Since Recon2.2 incorporates GAM ATP hydrolysis into the formation reaction for the protein biomass component, we do the same here. We add the associated substrates to the protein formation component, and we add the 4.37e-2 mass fraction to the existing specified mass fraction for protein, 7.21e-1, for a final protein mass fraction of 7.647e-1. We set the “Resolved small molecules” from “Table S1”^13^ as the “other” biomass component. Next, for each biomass component formation reaction, if equation (AA-7) (Supplementary Information and Results) does not hold, substrate coefficients were re-scaled to do so. For the carbohydrate biomass component specifically, the molecular weight of glycogen specified in “Table S1” is less than that of a glucose monomer; so, we use the Recon2.2 assigned glycogen molecular weight while re-scaling.

In Richelle et al.^50^, exchange reaction bounds were specified using measurements reported in “Table S2”. Exchange reactions that were removed during mCADRE model-extraction but experimentally reported in this table were not retained. Here, we added those reactions back, assuming that if they were experimentally measured, they were present in the cell line despite results of the model extraction step. Additionally, in instances where the M-Model forced flux through reactions, we relaxed the boundaries. In other words, if the lower bound was ≥ 0, we set it to 0, and if the upper bound was ≤ 0, we set it to 0. This modification to flux bounds was applied to exchange reactions as well, which somewhat deviates from *in vivo* accuracy since these bounds were set according to experimental measurements as reported in “Table S2” of Richelle et al.^50^, but was necessary for model feasibility.

### Construction and Growth Analysis of Multiple Cell Line ME-Models

To evaluate the broader applicability of the humanME framework beyond K-562, we constructed corresponding ME-Models for 15 additional cancer cell lines using context-specific Recon2.2-derived M-Models adapted from Richelle et al.^50^ as inputs. Each input model was preprocessed and reformatted as described above, including biomass reaction re-formatting, restoration of experimentally measured exchange reactions removed during context extraction, and inclusion of the original demand reactions in the M-Models. For each cell line, the same gold-standard PSIM and ME-Model building pipeline used for the K-562 analysis were applied to generate cell line-specific expression, transport, degradation, complex formation, and coupling reactions. For each cell line, the corresponding ME-Model was solved using qMINOS in quad precision and the maximal feasible growth rate was identified by binary search, as described above for the K-562 model. Of the 44 possible Recon2.2-derived M-Models, 16 were feasible.

As discussed in the Supplementary Material, the additional machinery constraints often over-constrain the ME-Model, leading to infeasibility or low growth. Identifying specific reaction bottlenecks in each cell line as done for K-562 becomes unwieldy. Thus, for comparison to corresponding M-models, across cell lines, we relaxed nutrient exchange boundaries 2-fold, corresponding conceptually to conditions in which nutrients-limitation is less substantial. Relaxing the bounds yielded one additional feasible model. In parallel, the corresponding input M-Model was solved using the same biomass objective and exchange reaction bounds to obtain the maximal FBA growth rate for direct comparison. To compare growth predictions across cell lines, we recorded the optimal biomass flux identified by each M-Model and its corresponding ME-Model with this relaxed exchange boundaries. Comparisons were performed on matched model pairs only.

### ME-Model FBA and FVA

ME-Model FBA solutions were post-processed to correct numerical precision artifacts arising from solver feasibility tolerances. Specifically, flux values that violated reaction bound constraints were set to their bound. To compare M-Model fluxes with ME-Model fluxes from the metabolic module, we mapped reactions back from the ME-Model to the original input M-Model. During ME-Model building, isozymes (reactions with an “OR” in the GPR) are each split into a separate reaction. Additionally, reversible reactions are split into their forward and reverse directions. For FBA, reactions that were split due to isozymes were aggregated by taking the sum of their fluxes. Next, reversible reactions that were split into their forward and reverse directions were aggregated by subtracting the flux for the reaction in the reverse direction from that in the forward direction.

For FVA, first, upper and lower flux values were re-oriented for directional consistency in instances that a reaction represented the reverse direction. Next, ME-model reactions corresponding to the same M-model reaction were aggregated by taking the maximum upper flux value and minimum lower flux value across reactions. FVA reaction bounds for the M-Model were identified using cobrapy’s ‘flux_variability_analysis’ function. FVA reaction bounds of the ME-Model were found by iterating through each reaction and maximizing and minimizing the flux through that reaction.

To compare model efficiency, all optimizations were performed with the biomass reaction as the objective, and the biomass flux was constrained to the maximum growth rate predicted by the ME-model. This ensured that differences in total absolute flux were not attributable to differences in biomass production. FBA solutions for both the M-model and ME-model were computed using the qMINOS solver in quad precision^75^. M-model solutions excluding thermodynamically infeasible loops were obtained by post-processing the FBA solution with the CycleFreeFlux algorithm^50^ using cobrapy’s ‘loopless_solution’ function. M-Model parsimonious FBA solutions were computed using cobrapy’s ‘pfba’ function. The M-model solution space was characterized by sampling 1000 feasible flux distributions using cobrapy’s ‘sample’ function. All *cobrapy*^*43*^ (v0.18.1) functions were run using default parameters.

To quantify the relative reliance on glycolysis versus oxidative phosphorylation, we computed the ATP flux ratio (AFR), defined as the ratio of glycolytic to oxidative phosphorylation (OxPhos) ATP-producing reaction fluxes, as previously described^57^. We additionally computed a modified AFR that accounts for ATP-consuming reactions by subtracting ATP-consuming from ATP-producing reaction fluxes within each pathway prior to ratio calculation; since we are aggregating by subtraction, we use total fluxes rather than mean fluxes.

### Transcriptional Analysis

In the ME-Model, transcription is represented by a final “synthesis” reaction that produces the mRNA species subsequently translated. For genes with synthesis reactions in multiple compartments, multiple such reactions may exist. Gene-level mRNA expression fluxes were therefore computed by summing fluxes across all corresponding synthesis reactions in the FBA solution.

log1p(TPM) K-562 RNA-sequencing data was downloaded from DepMap (version: Public 25Q3; file name: OmicsExpressionTPMLogp1HumanAllGenes.csv). Gene symbols were mapped to HGNC identifiers for consistency with the ME-Model, yielding 1,047 mapped genes out of 19,215 measured. For correlation analyses, 1,765 genes intersected between the RNA-sequencing dataset and the 1,778 genes represented in the K-562 ME-Model. Null distributions were generated by permuting gene labels within this intersecting set 10,000 times. For both growth and electron transport chain objectives, FBA solutions were computed at 95%, 97.5%, 99%, and 100% of the optimal objective value to account for potential objective trade-offs and hedging and to avoid edge cases at strict optimality. Correlation analyses were performed across these solutions, and the maximally correlated solution is reported. The ETC objective was set as maximization of the sum of fluxes through Complexes I-IV.

Enrichment analysis was performed using Metascape^77^, with the background defined as the intersecting gene set and the target list defined as “zero-flux” genes with log1p(TPM) ≥ 2 and transcriptional flux equal to zero. Protein– protein interaction–based enrichment was used because the ME-Model gene set, derived from Recon2.2, is biased toward metabolic genes and incompletely covered by standard pathway annotations.

## Code and Data Availability

Data files needed for building the ME-Model can be found at https://doi.org/10.5281/zenodo.20453308. The package repository can be found at https://github.com/hmbaghdassarian/human_me and scripts for analyses can be found at https://github.com/hmbaghdassarian/me_analyses_1.

## Authors, Contributions, and Acknowledgements

H.M.B. and N.E.L. conceived the work. All authors provided important insights for formulating and analyzing the ME-Model. H.M.B. implemented the Python package and performed analyses of the K-562 cell line. P.D.G. and L.D. implemented code for constructing and analyzing ME-Models for multiple cell lines. J.T.B. and L.Y. provided input on deriving the coupling coefficients. J.T.B provided input on formulating the biomass reaction and resolving K-562 feasibility due to an incomplete lipid synthesis network. L.Y. provided input on implementing the solver. E.A. provided input on improving model feasibility and implemented code to parse reaction GPRs. S.G. assisted with model context-extraction. H.M.B. wrote the paper and all authors carefully reviewed, discussed and edited the paper. This work was done with support from NIH R35 GM119850 (NEL) and a grant from Sartorius Stedim.

