## Supplementary Information for "Human Genome-Scale Models of Metabolism and Gene Expression Reveal Resource Constraints of Cancer Cell Lines"

### Supplementary Material

#### Supplementary Figures

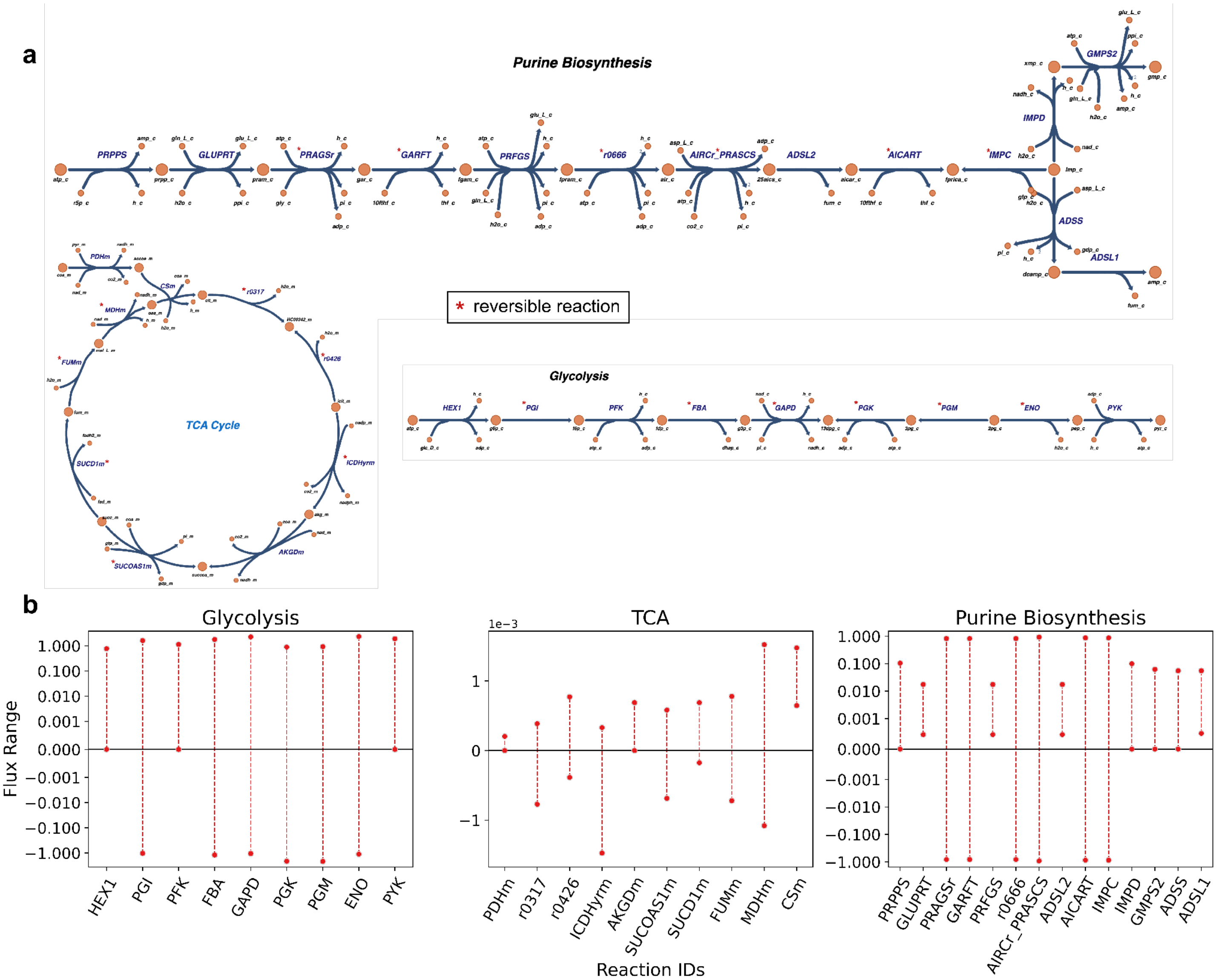

**Supplementary Figure 1: (a)** Escher map visualization of the three pathways analyzed by FVA. Reaction arrows point in the forward direction as specified by Recon2.2, with reversible reactions denoted by a red asterisk. **(b)** FVA of the ME-Model for reactions in glycolysis (left panel), the TCA cycle (middle panel), and purine biosynthesis (right panel). Individual reactions (x-axis) are plotted against minimum and maximum fluxes (y-axis, symlog scale) that maintain 95% of the maximal growth rate identified by each model. This is the same data as Fig. 2b, but only showing the ME-Model results to visualize the y-axis at higher resolution.

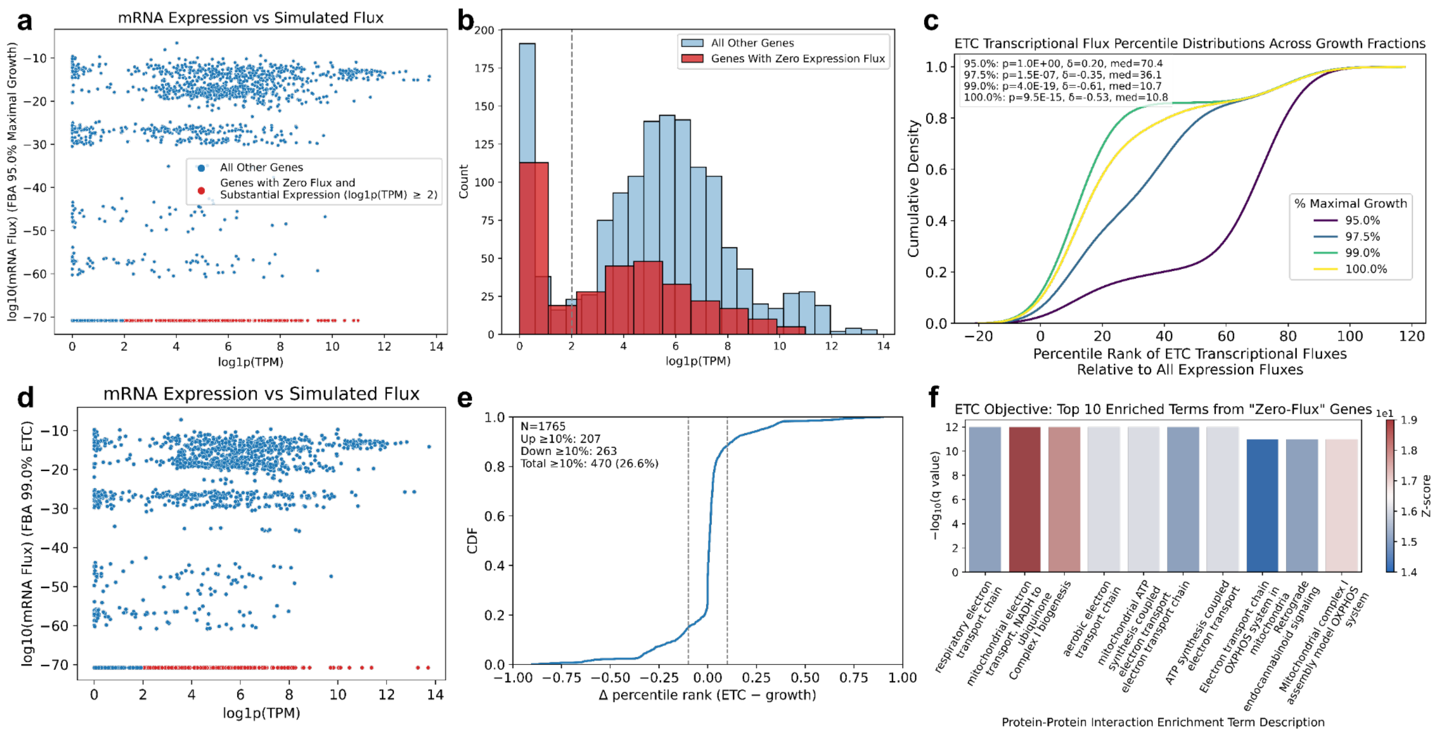

**Supplementary Figure 2:** (**a**) Scatterplot showing the relationship between transcript abundance (x-axis; log1p(TPM)) and transcriptional flux (y-axis, log-transformed) inferred from flux balance analysis (FBA) at 95% of maximal growth. This panel shows the same data as Fig. 3a prior to rank normalization and without kernel density estimates. Red points denote genes classified as "zero-flux" (log1p(TPM) ≥ 2 and FBA flux = 0), which are subsequently used as input to Metascape enrichment analysis. **(b)** Histogram of log1p(TPM) values for the 1,765 genes intersecting between the RNA-sequencing dataset and the K-562 ME-Model. Red bars indicate genes with zero FBA flux (regardless of expression level), while blue bars indicate genes with non-zero FBA flux. The gray dashed vertical line denotes the heuristic expression threshold used to label genes as "zero-flux." **(c)** Cumulative density plots of percentile-ranked transcriptional fluxes for monomeric subunits of Complexes I–IV across FBA simulations performed at 95–100% of the maximal growth rate, with growth as the objective. At each growth rate, plots are annotated with Mann–Whitney U test p-values and Cliff's delta effect sizes comparing ETC gene transcriptional fluxes to all other genes in the ME-Model. Median values indicate the median percentile rank of ETC genes relative to all other genes. **(d)** Same visualization as panel (a), but for the ETC objective at 99% of the maximal value. **(e)** Empirical cumulative distribution of objective-dependent rank shifts in transcriptional flux. Δ percentile rank is defined as the difference between percentile-ranked transcriptional fluxes under electron transport chain (ETC) and growth objectives (ETC − growth). Dashed vertical lines indicate the ±10 percentile-rank threshold used to define substantial rank shifts, with genes falling outside the interval representing substantial objective-dependent rank shifts. **(f)** Bar plots of the top 10 enriched terms for ETC objective "zero-flux" genes from Metascape's protein–protein interaction enrichment analysis, ordered by Benjamini–Hochberg FDR-corrected q-values and colored by z-score. This is analogous to Fig. 3b for the ETC objective.

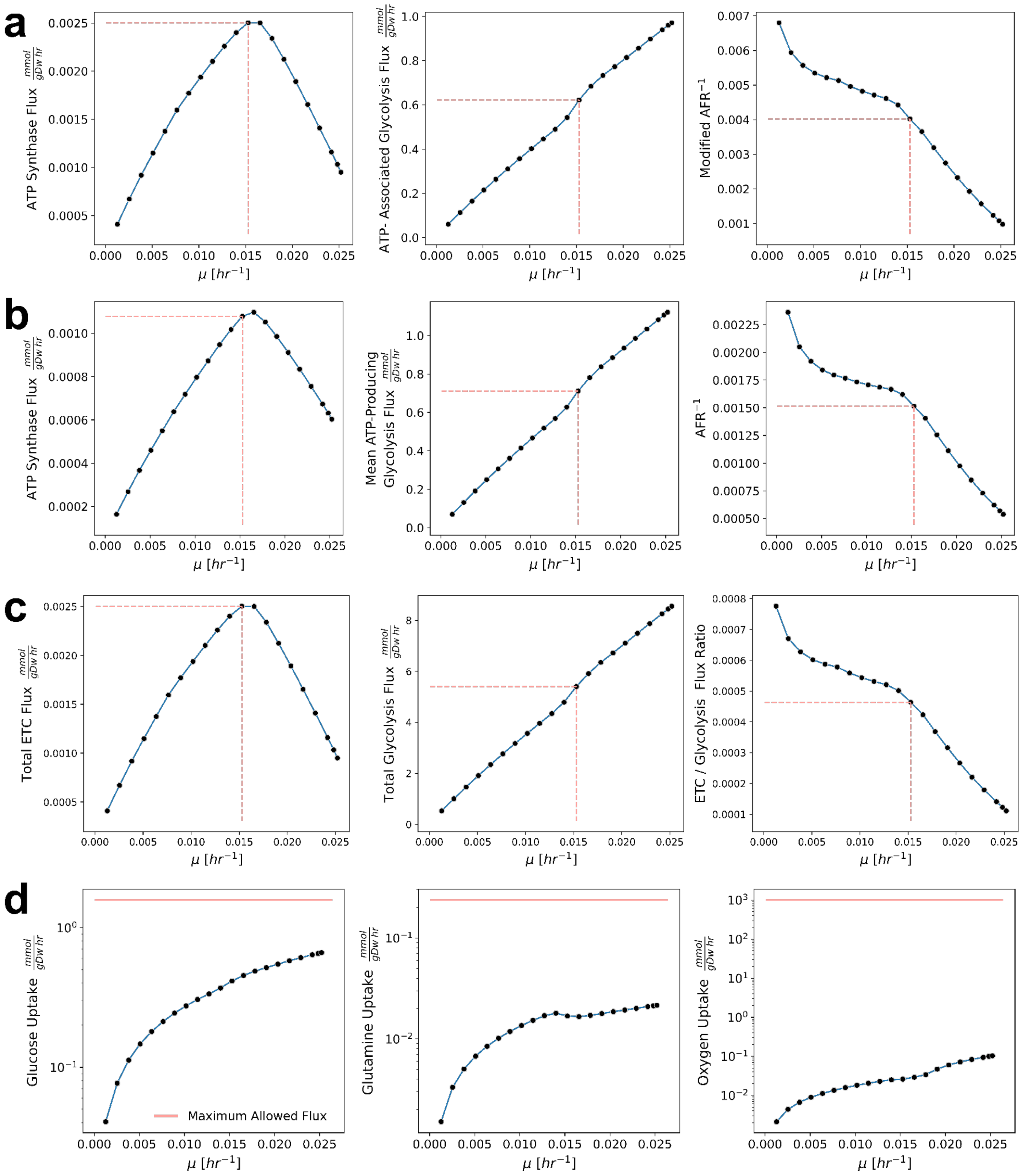

**Supplementary Figure 3:** All panels show FBA flux solutions with ETC as the objective at a given growth rate, and ETC enzymes at standard proteome cost (i.e., standard ME-Model formulation). Black points represent individual FBA solutions, while blue lines connect solutions across growth rates. Red dashed lines delineate the growth rate at which ETC is maximized (vertical line) as well as the corresponding pathway flux value (horizontal line). **(a)** Pathway fluxes were aggregated by subtracting the summed ATP-consuming reactions from summed ATP-producing reactions, yielding the inverse modified AFR (i.e., OxPhos to glycolysis ratio) shown in the right panel. This exactly matches the data in Fig. 3c of the main text, without the cost-free version displayed. **(b)** Mean pathway fluxes shown for ATP-producing reactions in oxidative phosphorylation (OxPhos) and glycolysis (left and middle panels). The ratio of OxPhos to glycolysis yields the inverse AFR (calculated as in the original reference[1]) shown in the right panel. **(c)** Total (sum of) pathway fluxes shown for ETC reactions (Complexes I-IV) representing oxidative phosphorylation (OxPhos) and glycolysis (left and middle panels). The ratio between the two pathways is shown on the right panel. **(d)** Exchange reaction uptake rates for glucose, glutamine, and oxygen. Solid red lines represent the upper bound constraint for uptake.

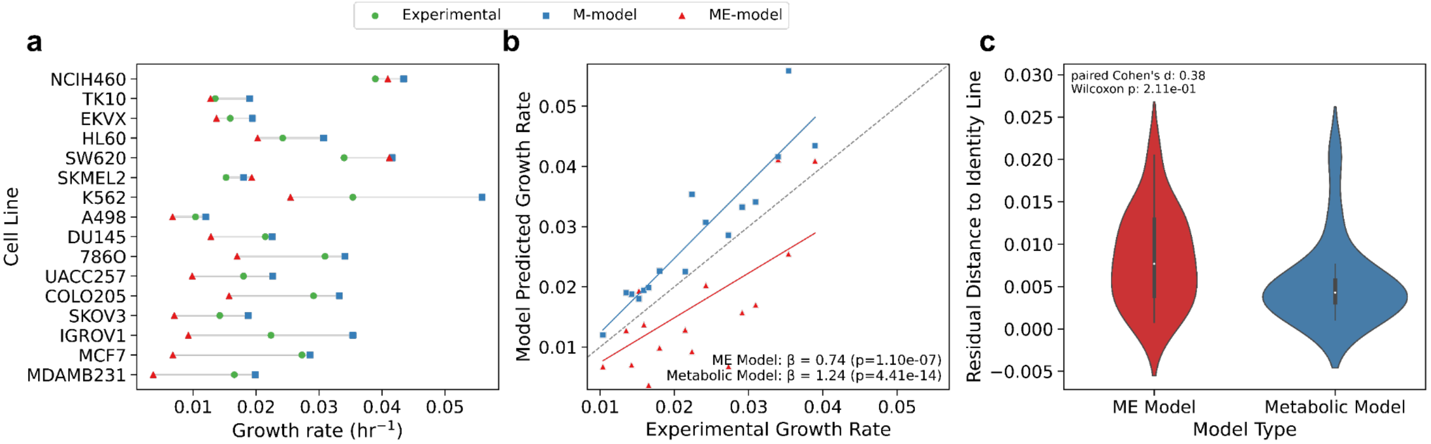

**Supplementary Fig. 4: ME- and M-Model growth rate prediction without exchange bound relaxation.** Panels are analogous to those of Fig. 2c-e when bounds are relaxed 2-fold. **(a)** Experimental growth rates alongside M- and ME-Model maximal growth rates across the 16 cell lines for which feasible matched ME-Model/M-model solutions were obtained. Cell lines are ordered by the most accurate ME-Model predictions. The ME-Model systematically predicts lower maximal growth rates than the paired M-model, consistent with the additional machinery constraints imposed by the expression module. **(b)** Scatter plots and OLS regression lines compare model predicted growth rates to experimental growth rates. OLS coefficients are annotated alongside their Wald test p-values. Coefficients were estimated using models without an intercept to enable direct comparison to the identity line y=x (gray), with coefficients closer to 1 indicating more accurate predictions across cell lines. **(c)** Violin plots summarize the absolute distance of each scatter point in panel (d) to the identity line , representing deviation from perfect prediction.

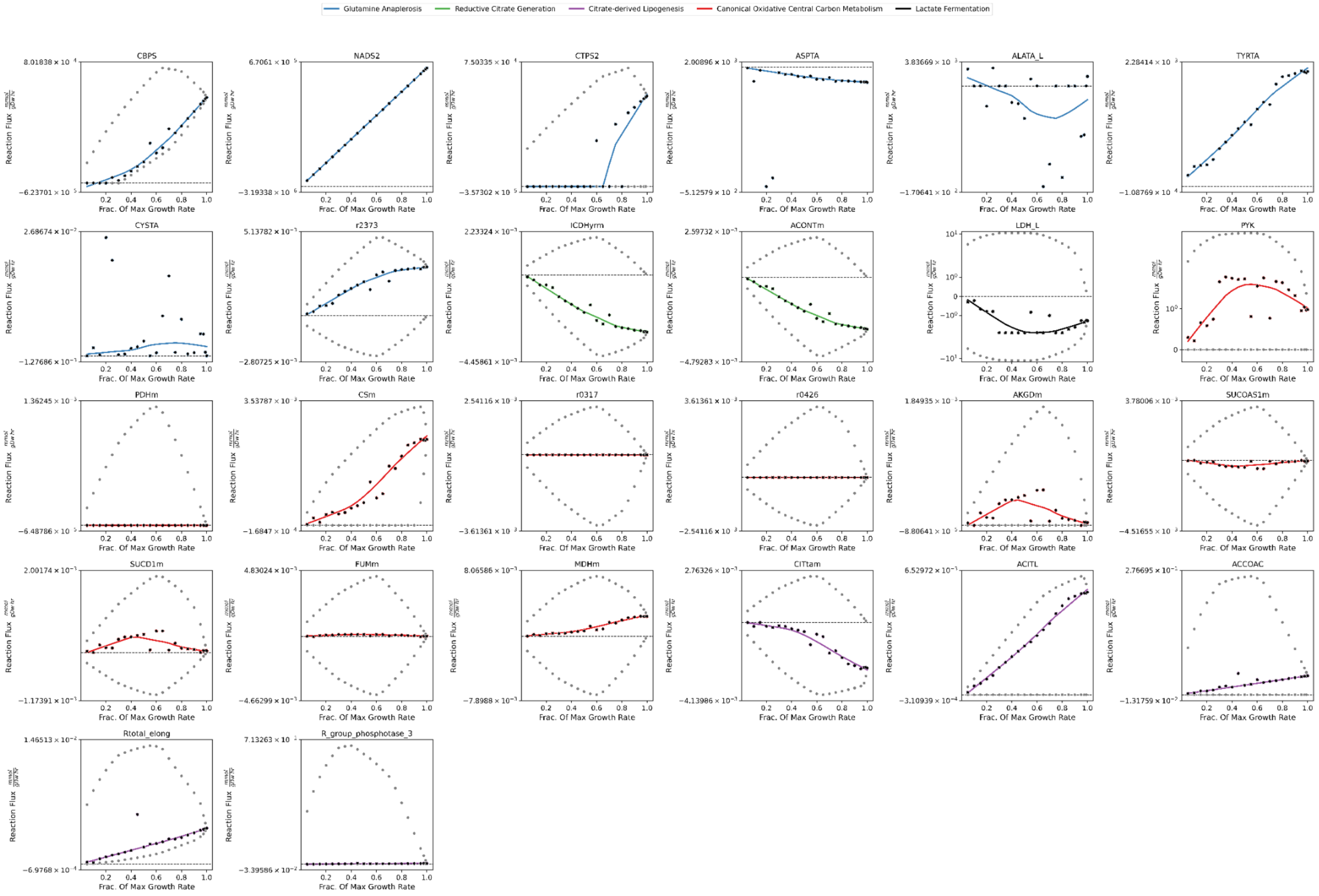
Glutamine Anaplerosis
Reductive Citrate Generation
Citrate-derived Lipogenesis
Canonical Oxidative Central Carbon Metabolism
Lactate Fermentation
CBPS
NADS2
CTPS2
ASPTA
ALATA_L
TYRTA
CYSTA
r2373
ICDHyrm
ACONTm
LDH_L
PYK
PDHm
CSm
r0317
r0426
AKGDm
SUCOAS1m
SUCD1m
FUMm
MDHm
CITtam
ACITL
ACCOAC
Rtotal_elong
R_group_phosphotase_3
Reaction Flux
Frac. Of Max Growth Rate

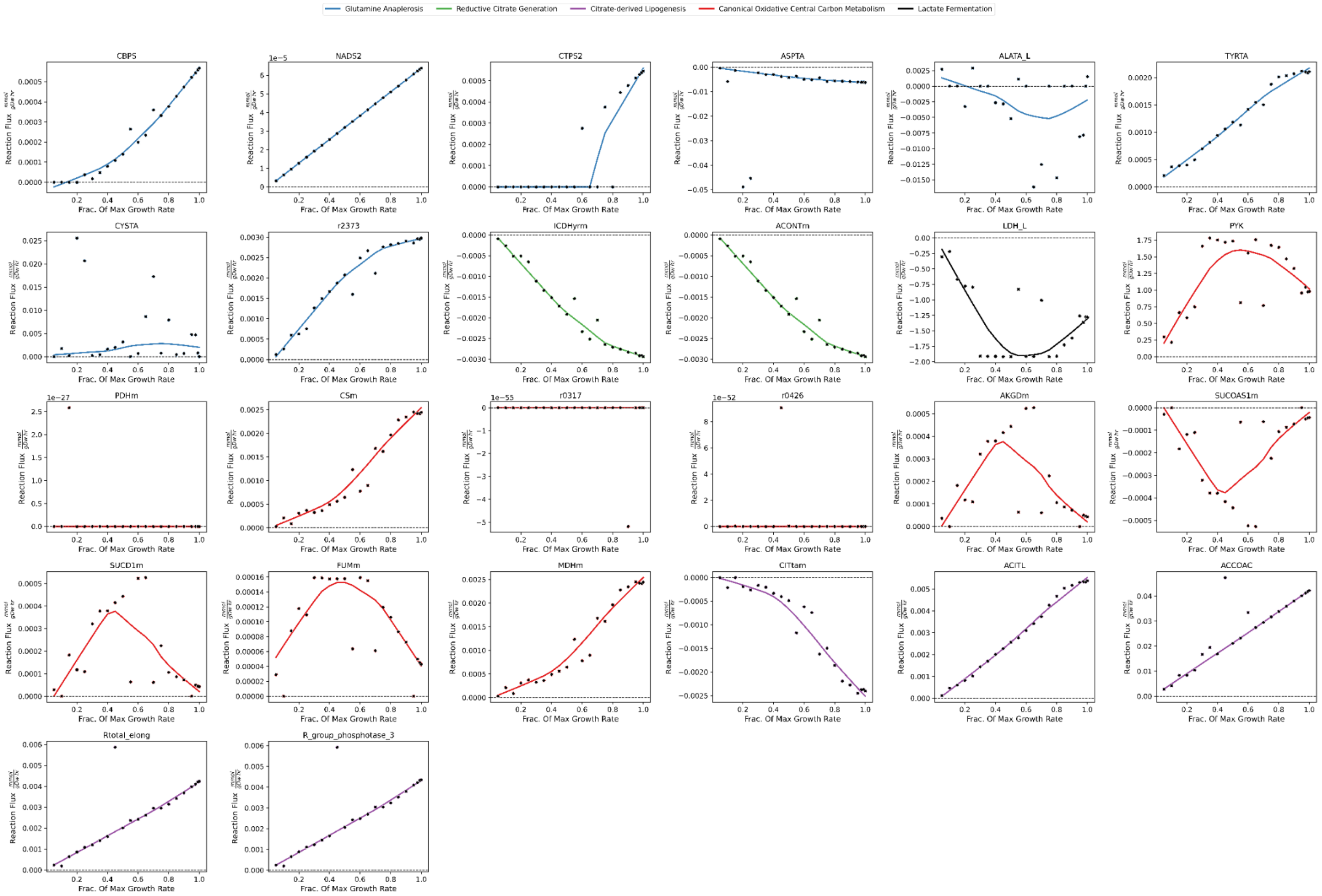

**Supplementary Fig. 5:** *Individual reaction fluxes as a function of growth rate identified by FBA and select FVA solutions under a growth objective. All reactions from various metabolic processes associated with citrate metabolism and lipogenesis are visualized. The x-axis is normalized to the maximum ME-Model growth. Gray scatter points visualize FVA bounds, black scatter points visualize individual FBA solutions, and lines visualize loess regression curves of the FBA solution. FBA solutions are also visualized in the absence of FVA ranges to emphasize trends that may be lost due to scale differences relative to FVA bounds.*

### Supplementary Methods

#### Building the ME Model

##### Gene Product Sequences and the Protein-Specific Information Matrix

We construct a gold-standard Protein-Specific Information Matrix (PSIM) containing all the gene features (Table 1) needed for the expression of metabolic module machinery (i.e., genes in Recon2.2) and the expression module machinery. While we include other genes (i.e., for expression of non-machinery gene products), we only ensure that the features for machinery genes are correctly encoded (e.g., macromolecular sequences contain only identified and canonical residues, there are not sequence mismatches between pre-mRNA, mRNA, and protein, etc.). For sequence features, cDNA and protein sequences were taken from MANE Select[2]. In instances where MANE did not contain information for a given gene, Refseq Select[3] was used in its place. Refseq Select is a superset of MANE Select. Refseq and MANE Select together yield consensus-isoform protein and transcript sequences for 18,495 unique genes. Of the 2,353 genes in the metabolic and expression module of the ME-Model, 78 were not covered by these databases. For these remaining 78, we use APPRIS[4]. We choose the isoform with the highest "PRINCIPAL" rating. The number of exons and gene sequences were downloaded from the ENSEMBL rest API. pre-mRNA and mRNA sequences were obtained by transcribing the gene and cDNA sequences, respectively.

Details on other gene features referred to in Table 1 are provided in corresponding subsections within “Building the ME-Model” in which those features are used. Additional formatting information on the PSIM can be found at the link: https://hmbaghdassarian.github.io/human_me/psim/. Users can provide a custom PSIM instead of using the gold-standard PSIM, in which case the gold-standard is only used for incorrect or missing values.

We also include additional information in the gold-standard PSIM that is not used for building the ME-Model. This includes a “Machinery” column indicating whether a protein is considered machinery according to the full Recon2.2 ('Metabolic'), the GPRs for expression reactions ('Expression'), both ('Both'), or neither ('Non-Machinery') and “Source” indicating which database the isoform sequences were attained.

**Table 1**: Gene features in the PSIM

| Name | Description | Default Value | PSIM-Gold Source |
|---|---|---|---|
| HGNC_ID[a] | The HGNC gene ID. There should be an entry for all genes that are included in the input M-Model GPRs and any expressed non-machinery. | | |
| PREMRNA_SEQ[b] | The gene pre-mRNA sequence. | The gold-standard PSIM values. Requirements include that values can only include 'A', 'C', 'G', 'U', the sequence length must be >= mrna sequence length, and the sequence length must be >= 3*protein sequence length. | [2–4] |
| MRNA_SEQ[b] | The gene mRNA sequence (isoform specific). | The pipeline will fill incorrect values with the gold-standard PSIM values. | [2–4] |
| PROTEIN_SEQ[b] | The gene protein sequence (isoform specific). | The pipeline will fill incorrect values with the gold-standard PSIM values. Requirements include values can only include one-letter amino-acid codes and the sequence length <= (mrna sequence length/3) | [2–4] |
| POLYA_LENGTH[c] | The length of the mature mRNA poly(A) tail. | If not provided, randomly draws from a fit Johnson SU distribution. | [5] |
| N_EXONS[c] | The number of exons in the premrna (isoform specific). Use to estimate the number of introns (as # of exons - 1). | 1 + <pre-mRNA sequence length>/6700 | [6] |
| TMD[d] | The number of transmembrane domains contained in the sequence. | 0 | [7] |
| DSB[d] | The number of disulfide bonds in the protein. | 0 | [7] |
| GPI[d] | Whether a GPI anchor is present in the protein. 0 if not present, 1 otherwise | 0 | [7] |
| OG[d] | The number of utilized O-linked glycosylation sites in the protein. | 0 | [7] |
| ALPHA_M[c] | The mRNA degradation rate ($hrs^{-1}$). | 0.06 $hrs^{-1}$ (median value from PSIM-Gold Source) | [8] |
| ALPHA_P[c] | The protein degradation rate ($hrs^{-1}$). | 0.02 $hrs^{-1}$ (median value from PSIM-Gold Source) | [9,10] |

| **Table 1**: Gene features in the PSIM | | | |
|---|---|---|---|
| **Name** | **Description** | **Default Value** | **PSIM-Gold Source** |
| PTR[c] | A gene-specific, tissue-independent protein to RNA ratio. | 65,163 (median value from PSIM-Gold Source) | [11] |
| LOCATION | The final model compartment of the protein. Required for non-machinery, disregarded for machinery (pipeline infers location from the corresponding reaction's compartments). | | |

a: default values unavailable, must be provided by user
b: default values unavailable, but if not provided or incorrect, will fill in with the gold-standard PSIM values when available (otherwise will error out)
c: standard default values used when not provided
d: only used for proteins that will be processed via the secretory pathway (ER, Golgi, extracellular membrane, plasma membrane, lysosome).

#### Preprocessing Inputs

Two inputs to the ME-Model building are preprocessed. The first is an M-Model obtained from a context-extraction of Recon2.2[12], which is a required input. The second is the PSIM, which defaults to the gold-standard if not provided by the user.

For the M-Model, first we check whether any reaction GPRs have the same enzyme repeated more than once. If they do, we remove them. In Recon2.2, this is the case for the following reactions: ACCOAC, OIVD1m, OIVD2m, OIVD3m, PFK, PI5P3K. We also ensure that exchange reactions are formatted as in Recon2.2, meaning that they are spontaneous and introduce metabolites to the model in two steps: 1) bounded input flux into the boundary compartment, and 2) an unbounded, reversible flux from the boundary compartment to the extracellular matrix. If present, we remove the gene "HGNC:4686", which is involved in catalysis of the reaction "GUACYC", because it is a pseudogene with no affiliated protein sequence. If present, we reformat genes with the format "HGNC:HGNC:#" to "HGNC:#". In Recon2.2, genes formatted this way are HGNC:HGNC:987 and HGNC:HGNC:2898. Next, there is a set of required metabolites for generating the gene expression reactions. If the M-Model does not contain these metabolites, first, we check whether it is present in another compartment and if it is, we add a transport reaction. If it's not present at all in the M-Model, we add it to the model using a sink reaction. Note that this results in the metabolite being generated at no resource cost to the model. We also add transport of hydrogen between the nucleus and cytosol, which is not present in Recon2.2. Finally, since the biomass reaction has to be reformatted (see Methods - Formatting the Biomass Reaction), we remove the biomass reaction as well as the biomass component formation reactions.

For a user-provided PSIM, preprocessing will ensure that the gene sequence features are present. The gold-standard PSIM is used to fill in missing values as well as any missing machinery genes. For other features, default values (Table 1) are used to fill in missing values.

#### tRNA Expression

Currently, tRNA reactions and molecules are constant. However, we note that the humanME codebase was written to accommodate custom mature tRNA, 5' leader 3' trailer, and intronic sequences[13]. The consensus tRNA sequence length was set to 72 bps for the mature tRNA, 6 bps for the 5' leader, and 9 bps for the 3' trailer, the most frequent values from "Table S4" of ref[14]. From this table, a position-independent consensus sequence was obtained from the frequency of each base in its relative position (i.e., normalized to total sequence length).

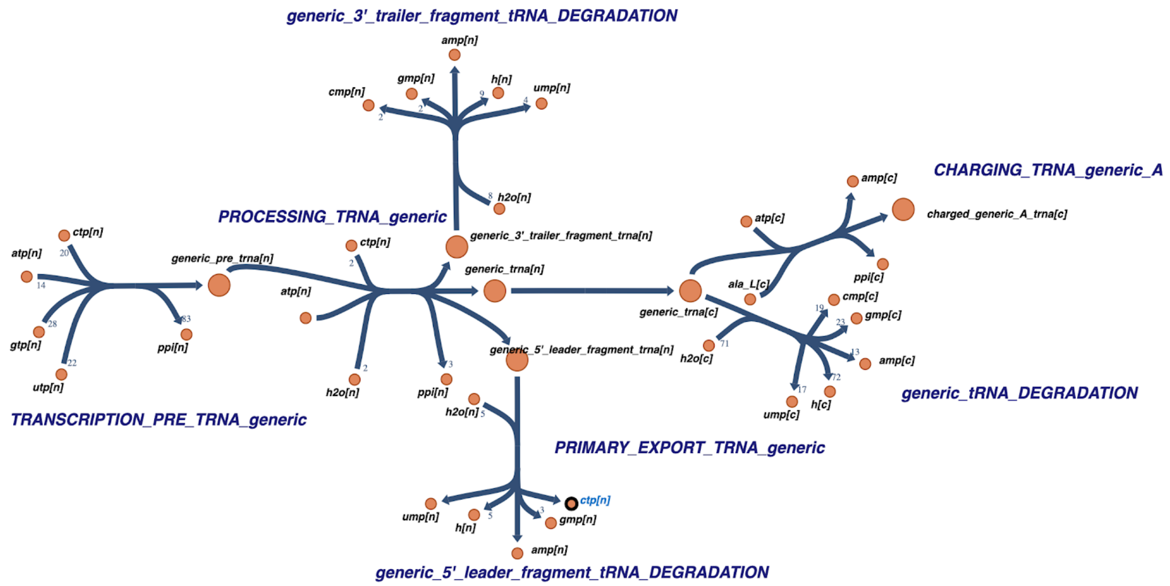

***Reaction network for mature tRNA synthesis.*** *Reactions for tRNA synthesis[15,16] include nuclear pre-tRNA synthesis by RNAP3[17], nuclear tRNA processing (e.g., 5' leader cleavage by RNase P[18], 3' trailer cleavage by RNase Z[19], addition of CCA[20], and splicing[21]), export to the cytosol[22], degradation of mature cytosolic tRNA[16], and tRNA charging[23].*

#### Ribosome Biogenesis

Ribosomal proteins are expressed[24] according to the gold-standard PSIM in the same manner as other gene products (see Methods - Gene Expression). RPL40 and RPS27A additionally include cleavage of ubiquitin by UCHL3[25].
rRNA sequences are obtained from NCBI. rRNA endonucleolytic cut sites are identified from ref[26,27] and scaled according to the NCBI sequence lengths. 5S rRNA is transcribed[28], processed[24,29], complexed with RPL5 and RPL11, and exported to the nucleus[30]. 18s, 5.8S, and 47S rRNA expression[26,27,29] includes 47S transcription[31] processing of intermediate pre-rRNAs.

pre-RNA processing occurs alongside formation of ribonucleoprotein complexes[29,32,3327,29,32–35] and nucleocytosolic export[36], ultimately resulting in 60S and 40S subunits that are joined[37] to form an active ribosome. Complex formation steps are irreversible, but the final active ribosome in the cytosol can dissociate into its individual subunits. rRNAs can be degraded in the cytosol by the exosome[38,39] upon ribosome complex dissociation.

#### Gene Expression: RNA

Gene expression reactions to produce mRNA and protein gene products associated with enzyme machinery (i.e., catalyze model reactions) are generated. First, nuclear pre-mRNAs are elongated from nuclear ribonucleoside triphosphates. Next, pre-mRNAs are processed, including addition of a 5' cap[40], 3' poly(A) tail[41,42], and splicing.

poly(A) tail lengths were obtained from the HeLA and iPSC organoid values in "Dataset 3" of ref[5]. Taking the average between these datasets, we identified the optimal distribution fit amongst all those available in the Python `scipy` package. Specifically, we fit each distribution to the mean across model systems. Next, we assessed goodness-of-fit using a Kolmogorov–Smirnov (KS) test comparing the empirical distribution of the data with the cumulative distribution function of the fitted model. The corresponding p-value were recorded for each distribution. We also quantified the discrepancy between the fitted probability density function (PDF) and the empirical density estimated from a 200-bin normalized histogram via the sum of squared errors (SSE). The fitted Johnson SU distribution demonstrated the best fit (lowest SSE and largest KS p-value, only distribution with p-value > 0.05) (Fig. MB). If the poly(A) tail length is not provided in the PSIM, it is randomly drawn from the fitted distribution. If it is provided, the poly(A) tail will be randomly drawn  from a normal distribution, with the mean as the provided value and the standard deviation calculated as described by an ordinary-least squares regression. Specifically, to get an expected standard deviation at a given length, we fit the regression predicting the standard deviation from the mean of poly(A) tails (both of these values are provided in "Dataset 3" for each gene). While the poly(A) tail is degraded over its lifetime, we assume it to be degraded in one single reaction.

The number of introns is used to calculate the total ATPs hydrolyzed during splicing, at a rate of 10 ATP per intron[43]. Lariats are degraded with the number of phosphodiester hydrolyzation events equal to the number of introns. In instances where the number of exons is provided by the PSIM, the number of introns is estimated to be the number of exons - 1 because there is an average of 1 fewer introns than exons per gene[6,44]. Otherwise, we estimate the number of introns as a function of the pre-mRNA sequence length (1 intron per 6.7 kbp[6]).

Next, processed products are exported to the cytosol via TREX[45–48]. Since transcription, processing, and export reactions are linear, there is an option to merge them into one reaction to reduce the total number of reactions in the ME Model and reduce solving time. Finally, mRNA is degraded[49] to ribonucleoside monophosphates in the 5' to 3' direction, including decapping by Nudix[50]. We note that humanME codebase includes the 3' to 5' degradation mechanism as well[49], but defaults to the 5' to 3' direction to have a single degradation reaction to use in the coupling (see Methods - Reaction Coupling).

#### Gene Expression: Protein

Once cytosolic mRNA is produced, the protein expression reactions depend on the final location of the protein. Proteins are transported to the final location wherein the reaction occurs. If the reaction occurs across more than one compartment, one is selected by the following criteria: If the extracellular matrix is one of those compartments, the protein location is the plasma membrane. Otherwise, if the cytosol is one of those compartments, it is disregarded. Next, the compartment with the most metabolites is assigned as the final location. If there is a tie, one is randomly selected.

We categorize those proteins with final locations in the cytosol, nucleus, mitochondrial matrix, mitochondrial intermembrane space, and peroxisome as "Cytosolic Transport" (i.e., they are transported to those compartments directly from the cytosol[51]). We categorize those proteins with final locations in the endoplasmic reticulum (ER), Golgi apparatus, lysosome, plasma membrane, and extracellular matrix as "Secretory Transport" (i.e., they are transported to those compartments via the secretory pathway).

For proteins that undergo Cytosolic Transport, they are translated[52] in the cytosol. For proteins that undergo Secretory Transport, they undergo co-translational translocation. 1 GTP per amino acid is hydrolyzed during translation[52]. Proteins destined for the cytosol, peroxisome[53,54], and nucleus that are longer than 100 amino acids[55] undergo irreversible post-translational folding by HSP70 and HSP40[56–58] in the cytosol. Proteins destined for the mitochondria undergo folding during transport. They are first translocated to the mitochondrial matrix[59], and those destined for the intermembrane space are further transported via the OXA complex[60]. Proteins destined for the nucleus can either undergo passive diffusion (if < 40kDa[51]) or classical nuclear import[61].

Cytosolic proteins undergo degradation via the ubiquitin-proteasome pathway (Fig. MC), since this accounts for 70% of protein degradation[62]. Ubiquitin is expressed like other cytosolic proteins for UBC and UBB genes, without folding and converted to monomers using the deubiquitinase USP5[63]. Proteins targeted for degradation are polyubiquitinated by 4 monomers via the E1-E3 ligases. 1 ATP is consumed per monomer[64]. Next, proteins are degraded by the 26S proteasome[65], with 2 ATP hydrolysis events per amino acid[64]. This ATP hydrolysis rate is also assumed for all other degradation and translocation reactions unless otherwise specified. Proteins destined for the cytosol, peroxisome, ER and Golgi via retro-translocation and ERAD, and nuclear proteins that can undergo passive nucleocytoplasmic diffusion can be degraded by this mechanism. Nuclear proteins can otherwise be degraded by a nuclear proteasome. Mitochondrial matrix and intermembrane proteins are degraded by LON protease[66], with 2 ATP hydrolyzed per amino acid, and i-AAA protease[67], respectively. Peroxisomal proteins can also be degraded directly in the peroxisome via LONP2[53], with 2 ATP hydrolyzed per amino acid.

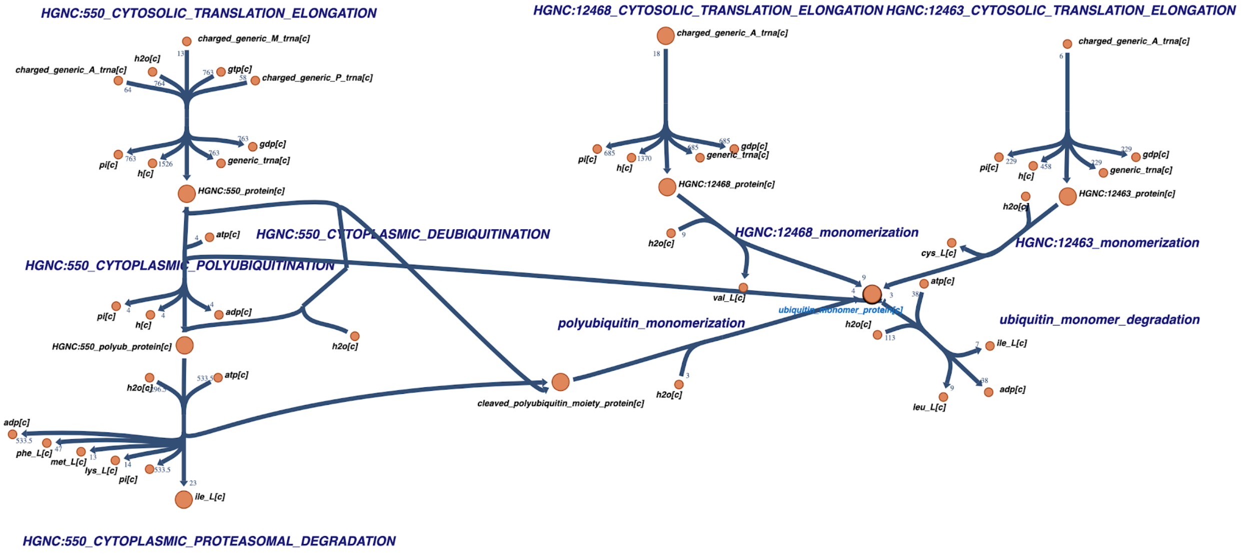

***Reaction network for cytosolic protein degradation.***

Transport reactions for proteins that undergo secretory transport are adapted from ref[7], with the addition of degradation reactions. Proteins that are destined for the Golgi and ER undergo degradation via ERAD[68,69]. They are irreversibly “misfolded”, retro-translocated, and then undergo cytosolic degradation. Plasma membrane proteins and lysosomal proteins undergo lysosomal degradation via cathepsins. Plasma membrane proteins are targeted to the lysosome by first undergoing ubiquitination[70] and then endocytosis[71].

Once transported to their final location, proteins that are subunits of a complex (i.e., GPRs that contain “AND” boolean logic) are joined via non-covalent interactions formed by a spontaneous reaction. The reversibility of complex formation reactions is a user-provided input, defaulting to being reversible. The reasoning behind making this reversible is for the ME-Model to be more resource efficient, as the monomeric subunits can then individually be degraded or, if they participate in other reactions as part of other enzymes, they can be re-used without having to be completely synthesized again. The individual subunits and the complexes have associated degradation reactions. For enzyme complexes, any parameters used in coupling (see Methods - Reaction Coupling) are taken as the median across all subunits in the complex. Since GPRs do not specify complex stoichiometry, complexes are assumed to form at a 1:1 ratio; however, the humanME codebase is written to accept specified subunit stoichiometries.

#### Reaction Coupling

After creating the gene expression reactions, we create demand for gene expression reaction fluxes by coupling expression to metabolism[72]. mRNA reactions must be coupled to protein reactions, and protein reactions must be coupled to metabolic reactions (Fig. 1d). Coupling constraints not only provide a way to link biological layers within the GEM framework, but they can also improve accuracy of models by explicitly linking dependent biological processes.

Take a toy example with reaction 1: $A + B \rightarrow C$, and reaction 2: $D \rightarrow E + F$. The flux through reaction 1, $v_1$, can be coupled to the flux through reaction 2, $v_2$ such that $v_1 = cv_2$, where $c$ is “coupling coefficient.” This can be done using one of two approaches. The first approach combines the two reactions with the coupling coefficient: $A + B + cD \rightarrow C + cE + cF$. The second approach uses one of the reaction products (or introduces a “proxy” metabolite product with no mass) in reaction 1 as a substrate in reaction 2, scaled by the coupling coefficient. Reaction 1 is $A + B \rightarrow C + P$, where P is the proxy metabolite. Reaction 2 is $cP + D \rightarrow E + F$. In humanME, we implement coupling constraints using the second approach.

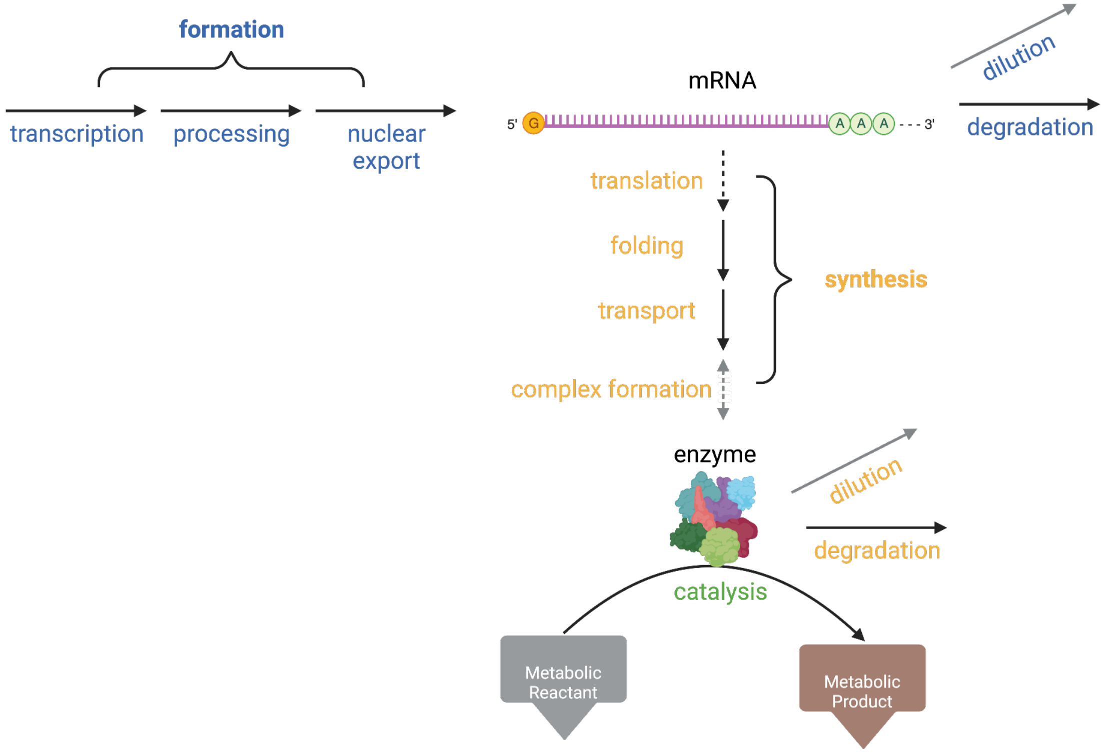

***The expression reaction network for a given gene in the ME-Model.*** *All labels are as described in Fig. 1. Brackets represent a series of typically linear reactions that are collapsed into a single reaction for simplification during coupling derivation. The gray dashed line indicates that complex formation only exists when the GPR contains "AND" boolean logic, otherwise the final monomeric protein product that is transported to its final location is the enzyme.*

To derive coupling coefficients, we first simplify the reaction network from that shown in the above diagram, which represents all expression reactions explicitly encoded by the model, to that shown in Fig. 1d, which merges linear series of reactions. This simplification allows for a simpler derivation of coupling constraints under steady-state. Variations of the reaction network visualized in Fig. 1d are commonly used to characterize genome-scale central dogma rates.[73–76]

Using this network, we derive two coupling coefficients that couple the mRNA layer to the protein layer (Fig. 1d). Specifically, $c_1$ couples mRNA formation to protein synthesis and $c_2$ couples mRNA degradation to protein synthesis. Similarly, we derive two coupling coefficients that couple the protein layer to the respective reactions that they catalyze. Specifically, $c_3$ couples enzyme formation to the catalyzed reaction and $c_4$ couples enzyme degradation to the catalyzed reaction. Note that $c_4$ is not implemented in prokaryotic ME-Models that assume growth rates are much larger than protein degradation rates[74,76,77]. Reaction GPRs are parsed to identify isozymes ("OR" boolean logic). During building of the ME-Model, each isozyme is separately expressed, and separately coupled to the reaction it is catalyzing. In other words, if a catalysis reaction is associated with two isozymes, the ME-Model will contain two versions of the catalysis reaction. Each version will contain the same metabolites and stoichiometry, except that they will be coupled to the respective isozyme with the isozyme-specific $c_3$ and $c_4$ coefficients.

Typically, the final reaction in the series of linear reactions is the reaction that is coupled. For example, if the active enzyme is a complex, the complex formation reaction is coupled to the catalyzed reaction in $c_3$; alternatively, if the enzyme is a monomer, the final transport reaction to the cell compartment is coupled to the catalyzed reaction in $c_3$,. The exception is in the protein synthesis reaction (denominator, Equations (11) and (12)) for $c_1$ and $c_2$. In this case, it is the first protein synthesis reaction (translation or co-translational translocation for Cytosolic and Secretory Transport enzymes, respectively) that is coupled to the final mRNA formation reaction. Logically, this makes sense because we are limiting the coupling to one branch point (i.e., a multi-localizing protein that is both cytosolically translated and co-translationally translocated).

For orphan reactions, i.e., those without a GPR, we assign a cytosolic dummy protein to catalyze them ("de-orphaning"). We reason that this is a more accurate representation of machinery resource costs than allowing the reaction to proceed spontaneously. The dummy protein represents a typical gene calculated from the features in the PSIM, limited to genes in Recon2.2. pre-mRNA and mRNA sequence lengths are calculated as the median across those in the PSIM. Protein sequence lengths are ⅓ this mRNA length. The base pair or amino acid in each position is assigned according to the relative frequency in that position as done for the consensus tRNA sequences (see Methods - tRNA Expression). Other features are calculated as the median of the values in the PSIM. Reactions that are not de-orphaned include: exchange reactions, demand reactions, expression module reactions that we curated and deliberately left without a GPR (e.g., complex formation), and some transport reactions. For transport reactions, we do not de-orphan those where all transported metabolites are less than 504 kDa[78] or if the reaction name contains "via diffusion".

**Table 2**: **Terms and notation for coupling constraints.**

| **Term/Notation** | **Definition** | **Units** |
| --- | --- | --- |
| [X] | Concentration of metabolite X | $mmol/gDW_{cell}$ |
| $i$ | mRNA of a specific gene | |
| $j$ | Protein of a specific gene | |
| $\tau_{1/2, i}$ | mRNA half-life | hr |
| $\alpha_{m,i}$ | First-order mRNA degradation constant<br>Calculated as $\ln(2)/\tau_{1/2, i}$ | $hr^{-1}$ |
| $\alpha_{p,j}$ | First-order protein degradation constant | $hr^{-1}$ |
| $k_{p,j}$ | First order rate constant for protein synthesis | $hr^{-1}$ |
| $MW_j$ | Molecular weight of protein $j$ | kDa |
| $SASA_j$ | Protein solvent accessible surface area.<br>Estimated as $MW_j^{0.75}$ | |
| $k_{cat,j}$ | Enzyme catalytic rate | $hr^{-1}$ |
| $PTR_{i,j}$ | The protein-to-rna ratio for a given gene. | |
| μ | Cell growth rate | $hr^{-1}$ |

We derive coupling constraints from first principles according to the simplified network above. They include a growth-dependent (dilution) and growth-independent (degradation) term. Growth-dependent terms account for dilution of molecules due to cell division. Since the biomass dilution reaction is a sink (see Methods - Formatting the Biomass Reaction for details), these components do not contribute to consumption and are not mass-balanced .

For $c_1$ and $c_2$, we derived coupling coefficients using context-independent parameters, i.e., those that we can expect to be fairly consistent across conditions. To this end, we utilize $PTR_{i,j}$, which, while being gene-specific, is demonstrated to be tissue-independent[79,80]. We also formulate coupling coefficients in terms of degradation rates rather than production rates, as degradation rates don't change a genes' distribution in Crick space, have a smaller dynamic range, and smaller contribution to the overall steady-state protein levels[74].

#### Coupling Coefficient Derivations

Coupling coefficients $c_1$ and $c_2$ are derived as follows:

From steady-state:

$$v_{\text{formation,i}} = v_{\text{dilution,i}} + v_{\text{degradation,i}} \quad (1)$$

$$v_{synthesis,j} = v_{dilution,j} + v_{degradation,j} \quad (2)$$

From first-order reaction laws:

$$v_{\text{dilution,i}} = \mu * [mRNA_i] \quad (3)$$

$$v_{\text{dilution,j}} = \mu * [protein_j] \quad (4)$$

$$v_{\text{degradation,i}} = \alpha_{m,i} * [mRNA_i] \quad (5)$$

$$v_{\text{degradation,j}} = \alpha_{p,j} * [protein_j] \quad (6)$$

Additionally:

$$PTR_{i,j} = \frac{[protein_j]}{[mrna_i]} \quad (7)$$

Substituting (4) and (6) into (2):

$$v_{synthesis,j} = (\alpha_{p,j} + \mu)[protein_j] \quad (8)$$

Substituting (7) into (8):

$$v_{synthesis,j} = (\alpha_{p,j} + \mu)PTR_{i,j}[mrna_i] \quad (9)$$

Substituting (3) and (5) into (1):

$$v_{formation,i} = (\alpha_{m,i} + \mu)[mrna_i] \quad (10)$$

Finally, if we define:

$$c_1 = \frac{v_{formation,i}}{v_{synthesis,j}} \quad (11)$$

$$c_2 = \frac{v_{degradation,i}}{v_{synthesis,j}} \quad (12)$$

Then, we can substitute (9) and (10) into the denominator and numerator of (11), respectively:

$$c_1 = \frac{(\alpha_{m,i} + \mu)[mRNA_i]}{(\alpha_{p,j} + \mu)PTR_{i,j}[mRNA_i]} = \frac{(\alpha_{m,i} + \mu)}{(\alpha_{p,j} + \mu)PTR_{i,j}} \quad (13)$$

And we can substitute (9) and (5) into the denominator and numerator of (12), respectively:

$$c_2 = \frac{\alpha_{m,i}[mRNA_i]}{(\alpha_{p,j} + \mu)PTR_{i,j}[mRNA_i]} = \frac{\alpha_{m,i}}{(\alpha_{p,j} + \mu)PTR_{i,j}} \quad (14)$$

Coupling coefficients $c_3$ and $c_4$ are derived as follows:

Assuming Michaelis-Menten kinetics:

$$v_{\text{catalysis,j}} = \frac{v_{\max}[S]}{K_{m,j} + S} = \frac{k_{\text{cat,j}}[protein_j][S]}{K_{m,j} + S} \quad (15)$$

assuming $K_m$ << [S]:

$$v_{\text{catalysis,j}} = k_{\text{cat,j}}[protein_j] \quad (16)$$

If we define:

$$c_3 = \frac{v_{\text{synthesis,j}}}{v_{\text{catalysis,j}}} \quad (17)$$

$$c_4 = \frac{v_{\mathrm{degradation,j}}}{v_{\mathrm{catalysis,j}}} \quad (18)$$

Then, we can substitute (8) and (16) into the numerator and denominator of (17), respectively:

$$c_3 = \frac{(\mu + \alpha_{\mathrm{p,j}})[protein_j]}{k_{\mathrm{cat,j}}[protein_j]} = \frac{(\mu + \alpha_{\mathrm{p,j}})}{k_{\mathrm{cat,j}}} \quad (19)$$

And we can substitute (6) and (16) into the denominator and numerator of (18), respectively:

$$c_4 = \frac{\alpha_{\mathrm{p,j}}[protein_j]}{k_{\mathrm{cat,j}}[protein_j]} = \frac{\alpha_{\mathrm{p,j}}}{k_{\mathrm{cat,j}}} (20)$$

#### Coupling Coefficient Parameter Values

The right-hand side of (13), (14), (19), and (20) are all parameters that are reported in the PSIM or can be calculated from the PSIM. $PTR_{i,j}$ is obtained from ref[11], and for genes that do not have a reported value, the median of 65,163 is used. $\alpha_{m,i}$ is obtained from ref[8], and for genes that do not have a reported value, the median of 0.061 $hr^{-1}$ is used. $\alpha_{p,j}$ is obtained from ref[9]. We took the median value across all cell lines. $\alpha_{p,j}$ was further supplemented with values reported from "Table S2" of ref[10]. Specifically, we include those categorized as short-lived in at least one cell line and not present in the first dataset. For genes that do not have a reported value, the median of 0.02 $hr^{-1}$ is used.

Since *in vitro* measurements of $k_{cat,j}$ are not widely available, we estimate the value from the protein solvent accessible surface area, $SASA_j$, as in COBRAme[81]. Specifically, we can set:

$$k_{cat,j} = SASA_j \frac{median(k_{cat})}{median(SASA)} \quad (21)$$

The median SASA is calculated across all unique metabolic enzymes (unique complexes or single-proteins). Across 5, 785 Recon2.2 enzymes, including complexes, the median SASA is 25.85. To get a median enzyme catalytic rate, we query the BRENDA database[82] for human enzymes. We disregard proteins reported as "mutants" in the comments. When a range was provided for a particular enzyme, the average was taken. Altogether, we retrieved 1638 catalytic rates from 238 unique EC-classes and 259 unique EC-class-protein pairs. For each EC-class-protein pair, we took the average across all reported values. We arrive at $median(k_{cat}) = 14338.8$ $hr^{-1}$.

#### Formatting the Biomass Reaction

Please first see the explanation and notation (Table 3) for the standard M-Model biomass reaction formulation (Supplementary Information and Results). The ME-Model adapts this formulation to enable variable RNA and protein mass fractions. Specifically, it directly modifies the M-Model biomass reaction as formatted in Supplementary Information and Results (equations AA-1 and AA-6).

In the ME-Model, each biomass component has a separate 1:1 input to the total biomass:

$$\text{for each } i \in n, \ \mathrm{biomass}_i \rightarrow \mathrm{biomass}_{\mathrm{tot}} \quad (22)$$

This 1:1 stoichiometry can be interpreted as follows: Each gram of the biomass component *i* produced contributes to 1 g of total biomass produced. This allows RNA and protein to be input to biomass at variable proportions. With this 1:1 ratio, using the same logic as equation (AA-4), the units of flux through equation (22) reactions simplify to $hr^{-1}$. Equation (22) has a lower bound of zero and an upper bound of 1000, but is constrained upstream by the growth rate μ as explained below.

Given the 1:1 ratio in equation (22), to enforce mass fraction of constant biomass components (those other than RNA and protein), each biomass component formation reaction must be reformatted from (AA-5) by scaling both reactants and products by $p_i$ as follows:

$$p_i s_{1,i} \mathrm{X}_{1,i} + p_i s_{2,i} \mathrm{X}_{2,i} + \ldots + p_i s_{m,i} \mathrm{X}_{m,i} \rightarrow p_i \mathrm{biomass}_i \quad (23)$$

The flux bounds of equation (23) are constrained to the growth rate (i.e., lower bound = μ and upper bound = μ). Since $biomass_{tot}$ directly determines growth rate by the biomass dilution reaction (see below for details), constraining the flux bounds of equation (23) by μ ensures that only $p_i$ counts of $biomass_i$ can contribute to $biomass_{tot}$ in equation (22) per unit flux. In other words, when the flux through equation (23) = μ, the production rate of $biomass_{i,}$ = $p_i\mu$ (see Table 3 definition of $v_j$). Note, the constraint equation (AA-7) to enforce mass balance of equation (AA-5) can be similarly written as equation (24) below to enforce mass balance through equation (23):

$$\sum_{j=1}^{m} \quad p_i s_{j,i} * MW(X_{j,i}) \qquad (24)$$
$$= p_i * \sum_{j=1}^{m} \quad s_{j,i} * MW(X_{j,i})$$
$$= p_i * 1 \qquad \text{(substitute in equation (AA-7))}$$
$$= p_i$$

For the variable biomass components (protein and RNA), instead of equation (23), biomass is produced (and consumed) directly in the gene expression reactions. For example, let's take a toy protein synthesis reaction that produces *k* units of $protein_A$, where *k* is the stoichiometric coefficient (Reaction A: amino acids → (*k*)$protein_A$). Considering the units of *k* [mmol $protein_A$/$gDW_{cell}$] (Table 3) and molecular weight [g $protein_A$/mmol $protein_A$] (Table 3), this can be converted to the amount of protein biomass produced by scaling *k* by the molecular weight of the protein [g protein/$gDW_{cell}$], matching the units of (AA-3):

$$k * MW(protein_A) = biomass_{protein} \quad (25)$$

Thus, we can add $biomass_{protein}$ as a product in Reaction A (amino acids → (*k*)$protein_A$ + (k)(MW($protein_A$))$biomass_{protein}$). Each reaction that forms or degrades RNA or protein, excluding coupled macromolecules (see Methods - Reaction Coupling section for reasoning), produces or consumes biomass in this manner. For RNA or protein consumption, the biomass term is in the substrates rather than the products. While there is an option to include an unmodeled fraction of the protein biomass as described in COBRAme[81], we set this to 0.

Finally, we can create a biomass dilution reaction that is analogous to the biomass consumption reaction in Supplementary Information and Results, with the only difference being that in the ME-Model, this is bounded by μ. This biomass dilution thus determines the growth rate, serving as the objective in the linear program (see Methods - Solving the ME-Model section for details).

We note that RNA and protein can only vary within their total mass fraction. Given that all other biomass components sum up to a mass fraction of *F*, where F < 1; then, while each of $p_{RNA}$ and $p_{protein}$ can vary, they are constrained by $p_{RNA} + p_{protein} = 1 - F$ (from equation AA-2).

Many biomass objectives include an ATP hydrolysis that represents GAM energetic costs[83]. In M-Models, including Recon2.2, the GAM ATP hydrolysis is often included in the protein biomass component formation reaction. Thus, replacing the (AA-5) format of the RNA and protein biomass components with equation (22) generally will eliminate the ATP hydrolysis for growth-associated maintenance (GAM). However, GAM is typically implemented in the protein biomass component formation reaction because protein synthesis costs represent a large portion of GAM costs and thus can be used as a proxy for GAM (see step 32 of ref.[83]). While there are other energetic costs which are not accounted for in the ME Model, e.g. error-checking and replication[84], we reason that removal of the explicit GAM ATP hydrolysis term by the ME-Model should not substantially underestimate the GAM costs, since the ME model explicitly accounts for the transcript and protein expression costs that constitute an overwhelming majority of GAM costs[43].

#### Solving the ME-Model

Given the large order of magnitude differences between standard stoichiometric coefficients and coupling coefficients, as well as expression ($10^{-16}$ - $10^{-8}$) and metabolic fluxes ($10^{-6}$ - $10^{3}$), we needed to use a high-precision solver. Thus, we implemented the qMINOS solver[85] through the QMINOS class in solveME[86]. Since some parameters in the ME-Model are a function of the variable μ (e.g., coupled metabolite coefficients and biomass reaction bounds), to create a Linear Program (LP), we must first substitute in a floating value for this variable. Once a value of μ is assigned, the model can be solved as in the M-Model; in other words, we can optimize for an objective of interest at a particular value of growth rate. In this case, if the objective of interest is growth rate, the solution will identify a flux equal to the assigned value. Thus, to maximize growth, we must identify the boundary at which the ME-Model becomes infeasible. To do this, we implement a binary search algorithm as described in COBRAme[81].

Finally, we implement an algorithm to maximize for an objective other than growth rate. To do so, we first need to identify the maximum growth rate μ*. Next, using growth rate values at a user-specified number of intervals within the feasible growth range [0, μ*], we can solve the LP and store the objective value. Finally, we identify the growth rate at which the objective value is maximized. Alternatively, we provide a higher resolution option to estimate the objective value as a function of μ using the scipy function “interp1D”. Specifically, we estimate the objective value across 1000 growth rate values specified at even intervals within the feasible growth range.

### Supplementary Information and Results

#### Additional Machinery Constraints Identify Missing Key Reactions in M-Model

When first constructing the ME-Model for the K-562 cell line, we observed that it was not feasible, even at very low growth rates (10 times less than experimental). While the M-model was feasible, the additional machinery constraints imposed by the ME-Model caused infeasibility, revealing key missing reactions that made the context-extracted K-562 metabolic network biologically implausible. This highlights one of the benefits of the ME-Model: identifying weaknesses in the model assumptions and constructing more biologically relevant metabolic networks.

Missing reactions necessary for feasibility in the ME-Model but not present in the initial context-extracted M-Model included those in the glycerol-3-phosphate shuttle for co-factor recycling, transport of a number of metabolites, exchange reactions (see Methods - Refining NCI-60 Cell Line M-Model Inputs for details of exchange reactions), and proline biosynthesis. More specifically for transport, we enabled transport of metabolites between the cytosol and the nucleus, as they were necessary in the nuclear compartment for various expression reactions: this included ribonucleotides, water, and diphosphate. Recon2.2 also does not contain a transport reaction for hydrogen between the cytosol and nucleus, so we added that. We also included transport of carbon dioxide and oxygen between the cytosol and extracellular matrix. Furthermore, we noticed that lipid anabolism was cyclical in Recon2.2 such that the metabolite “Rtotal_c”, used for downstream lipid biosynthesis, could not be produced. To resolve this, we added a proxy net reaction (accoa_c + 20 h_c + 7 malcoa_c + 14 nadph_c --> Rtotal_c + 7 co2_c + 8 coa_c + 6 h2o_c + 14 nadp_c) with the reaction ID “Rtotal_elong”.

Along these lines, only 16 of the 44 available cell lines yielded feasible ME-Models. Our troubleshooting of the K-562 model strongly suggests that this primarily reflects incompleteness of the input M-Model, as relaxing exchange bounds 10-fold recovered feasibility for only one additional cell line. Although the humanME preprocessing pipeline ensures availability of metabolites required for the expression module (e.g., through transport or sink reactions), it does not guarantee their complete downstream consumption for mass balance, which may contribute to infeasibility. Overall, these results highlight the importance of providing a high-quality input M-Model to humanME.

#### M-Model Biomass Reaction

Here, we outline how the M-Model biomass reaction is formulated, which is important for appropriately re-formatting it for the ME-Model (see Methods - Formatting the Biomass Reaction).

**Table 3**: **Term definitions for biomass.**

| Term/Notation | Definition | Units |
|---|---|---|
| $biomass_{tot}$ | The total biomass produced by the biomass reaction. 1g $biomass_{tot}$ = 1g cell dry weight ($DW_{cell}$) | |
| $biomass_i$ | Biomass component *i*<br>Typically: DNA, RNA, Protein, Lipid, Carbohydrate, Other | |
| $X_j$ | Metabolite *j* in the metabolic model | |
| *n* | The total number of biomass components | |
| *m* | The total number of metabolite precursors that form $biomass_i$ | |
| $MW_j$ | Metabolite $X_j$ molecular weight | kDa = g $X_j$/mmol $X_j$ |
| $p_i$ | Mass fraction of biomass component *i* | g $biomass_i$/g$DW_{cell}$ |
| $s_{j,i}$ | Molar-to-mass fraction of metabolite $X_j$ in biomass component *i*. | mmol $X_j$/g $biomass_i$ |
| $c^u_j$ | Stoichiometric coefficient of metabolite $X_j$ in the net biomass reaction (Equation AA-6).<br><br>Stoichiometric coefficients carry the units of concentration for the corresponding metabolite.<br><br>For each biomass component *i*, $c^u_{j,i} = p_i * s_{j,i}$ | mmol $X_j$/g$DW_{cell}$ |
| μ | Biomass reaction flux (i.e., cell growth rate) | 1/hr |
| $c^v_j$ | Stoichiometric coefficient of metabolite $X_j$ in a standard metabolic reaction. | mmol $X_j$/mmol (extent of reaction) |
| *v* | Metabolic reaction flux | mmol/g$DW_{cell}$/hr |
| $v_j$ | Metabolite $X_j$ consumption/production flux in a given reaction.<br><br>$c_j$*v in a metabolic reaction or $c^u_j$*μ | mmol $X_j$/g$DW_{cell}$/hr |

| | |
|---|---|
| | in the net biomass reaction (Equation AA-6). |

In this section, we explain two equivalent formulations of the biomass reaction (AA-6, or AA-1 combined with AA-5). Biomass reaction formulations ensure that one unit of reaction flux produces 1g $DW_{cell}$. To do so, units of stoichiometric coefficients (which hold the metabolite concentration units) and reaction flux differ from those of a standard metabolic reaction in the model.

The <u>biomass reaction</u> can be formulated as the sum of its components, each with a stoichiometric coefficient representing its mass fraction:

$$\text{For } i \in n,\ p_{i=1} * \text{biomass}_{i=1} + p_{i=2} * \text{biomass}_{i=2} + \ldots + p_{i=n} * \text{biomass}_{i=n} \rightarrow \text{biomass}_{\text{tot}} \quad \text{(AA-1)}$$

For mass balance, both the left- and right-hand sides of AA-1 must add up to 1g of total biomass. Given the definition of $p_i$ and this mass balance requirement, we have:

$$\sum_{i=1}^{n} p_i = 1 \quad \text{(AA-2)}$$

In addition to satisfying mass balance, the units of $p_i$ ensure that the production rate of $biomass_{tot}$ in AA-1 is equivalent to the growth rate. The production rate of $biomass_{tot}$ is equivalent to the consumption rate of all the biomass components. Bearing in mind that the production and consumption rates of metabolites are the corresponding stoichiometric coefficient multiplied by the flux (Table 3):

$$\mu = \text{biomass}_{\text{tot}} \text{ production rate} = \sum_{i=1}^{n} p_i * \mu \quad \text{(AA-3)}$$

From (AA-2) and (AA-3), the biomass reaction flux units are:

$$\sum_{i=1}^{n} \frac{g\ biomass_i}{g\ DW_{cell} * hr} = \frac{g\ biomass_{tot}}{g\ DW_{cell} * hr} = hr^{-1} \quad \text{(AA-4)}$$

Altogether, this reaction flux corresponds to the growth rate, and flux units and stoichiometric coefficient units ensure that 1g $DW_{cell}$ is produced per 1 unit of biomass reaction flux.

Each $biomass_i$ has its own <u>component formation reaction:</u>

$$\text{For } j \in m,\ s_{j=1,i}X_{j=1,i} + s_{j=2,i}X_{j=2,i} + \ldots + s_{j=m,i}X_{j=m,i} \rightarrow \text{biomass}_i \quad \text{(AA-5)}$$

Note, (AA-5) and (AA-1) are typically combined to create a <u>net biomass reaction</u> with stoichiometric coefficients $c^u_{j,i} = p_j * c_{j,i}$ (Table 3) as follows:

$$[(p_1 * s_{1,1})X_{1,1} + (p_1 * s_{2,1})X_{2,1} + \ldots + (p_1 * s_{m,1})X_{m,1}] + [(p_2 * s_{1,2})X_{1,2} + (p_2 * s_{2,2})X_{2,2} + \ldots + (p_2 * s_{m,2})X_{m,2})] + \ldots + [(p_n * s_{1,n})X_{1,n} + (p_n * s_{2,n})X_{2,n} + \ldots + (p_n * s_{m,n})X_{m,n})] \rightarrow \text{biomass}_{\text{tot}} \quad \text{(AA-6)}$$

For the ME-Model, the M-Model input should be formulated as (AA-1) and (AA-5), rather than the standard (AA-6). Units in the M-Model formulation as in (AA-6) follow such that the two formulations agree: To convert the net biomass reaction (AA-6) , the stoichiometric coefficients must be divided by $p_i$ to get the coefficients $s_{j,i}$ as in (AA-5). The stoichiometric coefficients ($c^u_{j,i} = p_i * s_{j,i}$) in (AA-6) are in [mmol $X_j$/g$DW_{cell}$] as delineated in Table 3. $p_i$ is in units of [g $biomass_i$/g$DW_{cell}$]. Thus, in (AA-5), the units of $s_{j,i}$ are derived from ($p_i * s_{j,i}$)/$p_i$, or [mmol $X_j$/g$DW_{cell}$] / [g $biomass_i$/g$DW_{cell}$], which gives [mmol $X_j$/g $biomass_i$]. So, if we multiply the coefficient $s_{j,i}$ by the molecular weight of the metabolite, we get units of [mmol $X_j$/(g $biomass_i$)] * [g $X_j$/mmol $X_j$] = [g $X_j$/g $biomass_i$]. Under these units, to achieve mass balance in (AA-5) analogous to the constraint (AA-2) that enforced mass balance in (AA-1), we have:

$$\sum_{j=1}^{m} s_{j,i} * \mathrm{MW}(\mathrm{X}_{j,i}) = 1 \quad \text{(AA-7)}$$

In other words, (AA-7) says that with the stoichiometric amounts $s_{j,i}$ of $X_{j,i}$ across all substrates in the biomass component formation reaction, (AA-5) produces 1g of $biomass_i$ per unit of reaction flux.

Additionally, following the same logic used in AA-1 - AA-4, the reaction flux units of AA-5 are also $hr^{-1}$:

- One unit of reaction flux through component formation must yield 1g of $biomass_i$.
- The total biomass metabolite precursor $X_j$ consumption rate is equal to the production rate of biomass: $biomass_i$ production rate = $\sum_{j=1}^{m} s_{j,i} * \mu$
- Units cancel out to produce those of growth rate: $\sum_{j=1}^{m} \frac{mmol\ X_i}{g\ biomass_i * hr} = \frac{g\ biomass_i}{g\ biomass_i * hr} = hr^{-1}$

Finally, there is a biomass consumption demand reaction to prevent $biomass_{tot}$ from accumulating, generating mass balance: "$biomass_{tot} \rightarrow$ ".
